# Interplay of BAF and MLL4 promotes cell type-specific enhancer activation

**DOI:** 10.1101/2020.06.24.168146

**Authors:** Young-Kwon Park, Ji-Eun Lee, Tommy O’Haren, Kaitlin McKernan, Zhijiang Yan, Weidong Wang, Weiqun Peng, Kai Ge

**Affiliations:** Adipocyte Biology and Gene Regulation Section, National Institute of Diabetes and Digestive and Kidney Diseases, National Institutes of Health (NIH), Bethesda, MD 20892; Laboratory of Genetics and Genomics, National Institute on Aging, NIH, Baltimore, MD 21224; Department of Physics and Department of Anatomy and Cell Biology, The George Washington University, Washington, DC 20052

## Abstract

Cell type-specific enhancers are activated by coordinated actions of lineage-determining transcription factors (LDTFs) and chromatin regulators. The SWI/SNF complex BAF and the histone H3K4 methyltransferase MLL4 (KMT2D) are both implicated in enhancer activation. However, the interplay between BAF and MLL4 in enhancer activation remains unclear. Using adipogenesis as a model system, we identified BAF as the major SWI/SNF complex that colocalizes with MLL4 and LDTFs on active enhancers and is required for cell differentiation. In contrast, the promoter enriched SWI/SNF complex PBAF is dispensable for adipogenesis. By depleting BAF subunits SMARCA4 (BRG1) and SMARCB1 (SNF5) as well as MLL4 in cells, we showed that BAF and MLL4 reciprocally regulate each other’s binding on active enhancers before and during adipogenesis. By focusing on enhancer activation by the adipogenic transcription factor C/EBPβ without inducing cell differentiation, we provide direct evidence for an interdependent relationship between BAF and MLL4 in activating cell type-specific enhancers. Together, these findings reveal a positive feedback between BAF and MLL4 in promoting LDTF-dependent activation of cell type-specific enhancers.

## Introduction

Throughout development, chromatin architecture undergoes dynamic changes that are critical for promoting appropriate cell type-specific enhancer activation and gene expression. These changes are coordinated by the action of transcription factors (TFs) and epigenomic regulators, including ATP-dependent chromatin remodeling complexes, which modulate chromatin accessibility and gene expression through the mobilization of nucleosomes^1,2^. The large multi-subunit SWItch/Sucrose Non-Fermentable (SWI/SNF) chromatin remodeling complexes have been shown to play a crucial role in cellular and tissue development^1^. SWI/SNF mutations are also frequently associated with human diseases, including over 20% of cancers and a variety of neurodevelopmental disorders^3,4^.

Mammalian SWI/SNF complexes exist in three different isoforms with distinct subunit compositions: BRG1/BRM-associated factor (BAF), polybromo-associated BAF (PBAF), and non-canonical GLTSCR1L-containing BAF (GBAF, also known as ncBAF)^5,6^. All three complexes contain an ATPase catalytic subunit, either SMARCA4 (BRG1) or SMARCA2 (BRM), and an initial core, composed of SMARCC1 (BAF155), SMARCC2 (BAF170), and SMARCD1/D2/D3 (BAF60A/B/C)^5,7^. The BAF complex contains SMARCB1 (SNF5, INI1, or BAF47), SMARCE1 (BAF57), and SS18 and BAF-specific subunits ARID1A/B (BAF250A/B) and DPF1/2/3 (Fig 1A). Like BAF, PBAF complex also consists of the initial core, SMARCB1 and SMARCE1 as well as PBAF-specific subunits PHF10, ARID2 (BAF200), BRD7, and PBRM1. The recently characterized GBAF is a smaller complex containing SMARCA4 or SMARCA2, the initial core, SS18, and GBAF-specific subunits GLTSCR1/L and BRD9^5-7^. These three distinct complexes exhibit different genomic localization patterns. BAF primarily binds on enhancers^8-10^. Conversely, PBAF is more localized on promoter regions, and GBAF binding sites are enriched with the CTCF motif^11,12^.

**Figure 1.**
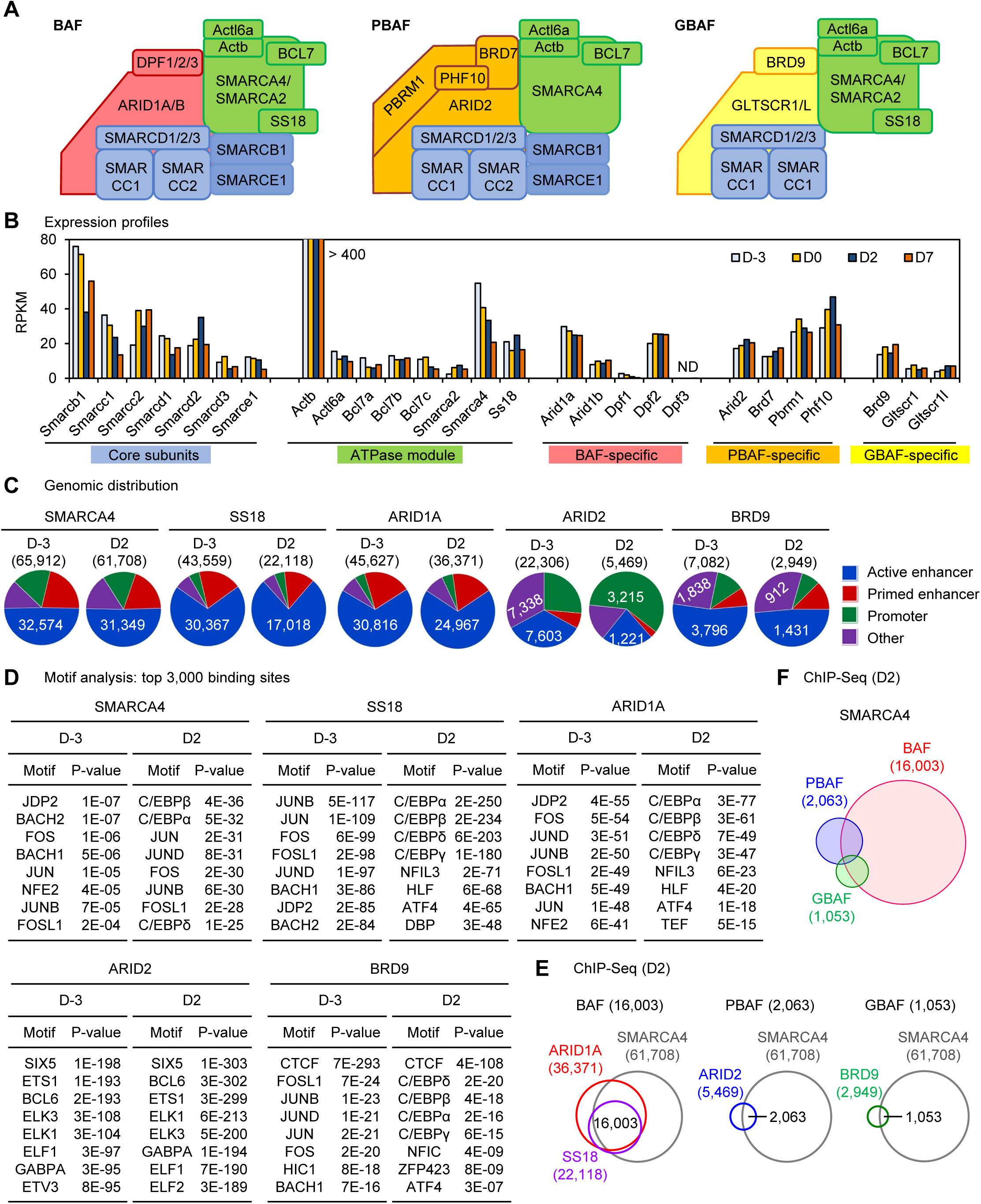
Expression and genomic binding of SWI/SNF subunits in adipogenesis. **(A)** Schematic of subunits in SWI/SNF complexes BAF, PBAF and GBAF (ncBAF). **(B)** Expression profiles of SWI/SNF subunits in adipogenesis. Published RNA-Seq data sets were used (GSE74189)^18^. D-3, day −3. D2, day 2. **(C)** Genomic distributions of SMARCA4 (BRG1), SS18, BAF-specific ARID1A, PBAF-specific ARID2 and GBAF-specific BRD9 before (D-3) and during (D2) adipogenesis were determined by ChIP-Seq. Promoters were defined as transcription start sites ± 1kb. Active enhancers were defined as H3K4me1^+^ H3K27ac^+^ promoter-distal regions. Primed enhancers were defined as H3K4me1^+^ H3K27ac^-^ promoter-distal regions. Numbers of binding sites are indicated. **(D)** Motif analysis of SWI/SNF subunits binding regions at D-3 and D2 of adipogenesis. Top 3,000 significant binding regions were used. **(E)** Venn diagrams depicting genomic binding of SMARCA4, SS18, ARID1A, ARID2 and BRD9 at D2 of adipogenesis. BAF, PBAF, and GBAF binding sites were defined as SMARCA4^+^ SS18^+^ ARID1A^+^, SMARCA4^+^ ARID2^+^, and SMARCA4^+^ BRD9^+^ regions, respectively. **(F)** Venn diagram depicting genomic binding of BAF, PBAF and GBAF at D2 of adipogenesis.

Enhancers are *cis*-acting regulatory regions containing TF binding sites and are responsible for facilitating cell type-specific gene expression through communication with promoters^13^. Enhancers are marked by H3K4 mono-methylation (H3K4me1), which is primarily placed by the partially redundant H3K4 methyltransferases MLL3 (KMT2C) and MLL4 (KMT2D)^14,15^. Along with H3K4me1, active enhancers are further marked with histone acetyltransferases CBP/p300-catalyzed H3K27 acetylation (H3K27ac)^16,17^. MLL3 and MLL4 are required for enhancer activation and cell type-specific gene expression during cell differentiation^14^. Specifically, MLL3/MLL4 controls enhancer activation by facilitating CBP/p300 binding on enhancers^18^. MLL3 and MLL4 are essential for embryonic development as well as the development of multiple tissues, including adipose and muscle^14,19,20^.

Of the three SWI/SNF complexes, BAF primarily interacts with enhancer regions. Recent studies have demonstrated that BAF is required for enhancer activation^8-10^. Following the deletion of BAF catalytic subunit SMARCA4 and core subunit SMARCB1, H3K27ac levels on enhancers are decreased, and enhancers fail to be activated. The deletion of SMARCB1 results in the destabilization of the BAF complex on the genome^9^. BAF-specific subunit ARID1A also plays a major role in BAF-mediated enhancer activation. Particularly, the loss of ARID1A has been shown to diminish BAF complex occupancy on enhancers^21^. SMARCA4 interacts with LDTFs including the adipogenic pioneer factor C/EBPβ and is required for adipogenesis^22-25^. Although the BAF complex plays an essential role in enhancer activation and development, many questions remain regarding how it interacts with other epigenomic regulators during enhancer activation.

In this study, we evaluated the interrelationship between BAF and MLL4 on cell type-specific enhancers using adipogenesis as a model system. Loss of SWI/SNF subunits in cells and mice provides evidence that BAF, but not PBAF, is critical for enhancer activation, adipogenesis and adipocyte gene expression. Given that MLL4 is similarly important for enhancer activation, cell differentiation and cell type-specific gene expression, we investigated the relationship between BAF and MLL4 before and during adipogenesis, revealing that BAF and MLL4 colocalize with LDTFs on active enhancers and facilitate enhancer activation in an interdependent manner. We further uncover the reciprocal regulation between BAF and MLL4 in adipogenic TF C/EBPβ-mediated enhancer activation.

## Results

### Expression and genomic binding of SWI/SNF subunits in adipogenesis

To explore the role of each SWI/SNF complex in cell differentiation and cell type-specific enhancer activation, we used differentiation of preadipocytes towards adipocytes (adipogenesis) as a model system. We first explored the expression profiles of SWI/SNF subunits at four key time points in adipogenesis: actively growing preadipocytes (day −3, D-3), over-confluent preadipocytes before the induction of differentiation (day 0, D0), immature adipocytes during differentiation (day 2, D2), and adipocytes after differentiation (day 7, D7)^18^. By RNA-Seq analysis, we found that all SWI/SNF subunits except *Dpf3*, a BAF-specific subunit, were expressed in adipogenesis. The expression levels of SWI/SNF subunits were largely invariant from D-3 to D7. Among the two enzymatic subunits, *Smarca4* was expressed at significantly higher levels than *Smarca2* throughout the differentiation (Fig 1B).

Next, we performed ChIP-Seq analyses to profile genomic localizations of BAF, PBAF, and GBAF complexes before (D-3) and during (D2) adipogenesis. We chose the enzymatic subunit SMARCA4, SS18, BAF-specific subunit ARID1A, PBAF-specific subunit ARID2 and GBAF-specific subunit BRD9. Each subunit exhibited a differentiation stage-specific genomic binding (Fig 1-S1A). ChIP-Seq identified distinct genomic distribution of each subunit in four types of regulatory elements: active enhancer (AE), primed enhancer, promoter and other regions defined as in our previous report^14^. SMARCA4 binding sites were distributed in all types of regulatory elements, but primarily on primed enhancers and AEs. While SS18 and ARID1A binding sites were mainly located on AEs, PBAF-specific ARID2 was strongly enriched on promoter regions especially during adipogenesis. GBAF-specific BRD9 binding sites were enriched on AEs and other regions (Fig 1C). By motif analysis of the top 3,000 binding sites of each subunit, we found that SMARCA4, SS18 and ARID1A binding regions were enriched with motifs of AP-1 family TFs Jdp2, JunD and Jun in preadipocytes (D-3), but with motifs of adipogenic TFs such as C/EBPα, C/EBPβ, and ATF4 during adipogenesis (D2). ARID2 and BRD9 binding regions were selectively enriched with motifs of ETS family TFs and CTCF, respectively (Fig 1D). These results demonstrate the distinct targeting of each complex in adipogenesis: BAF to enhancers, PBAF to promoters and GBAF to CTCF sites.

Further, we defined confident genomic binding regions of BAF (16,003), PBAF (2,063) and GBAF (1,053) at D2 of adipogenesis by selecting overlapping binding sites between SMARCA4 and complex-specific subunits (Fig 1E and Fig 1-S1B). Even though SS18 was reported as a common subunit in BAF and GBAF complexes, the majority of SS18 genomic binding sites (20,257/22,118) overlapped with those of BAF-specific ARID1A, and only a small fraction (791/22,118) overlapped with those of GBAF-specific BRD9. Among 17,165 SWI/SNF-associated SMARCA4 binding regions, BAF (16,003) exhibited substantially greater genomic occupancy than PBAF (2,063) and GBAF (1,053) during adipogenesis (Fig 1-S1B). Only a small subset (1,504/16,003, 9.4%) of BAF binding sites overlapped with those of PBAF or GBAF (Fig 1F). These findings suggest that among the three SWI/SNF complexes, BAF is the major regulator of enhancers in adipogenesis.

### BAF, but not PBAF, is required for adipogenesis

To address the functions of SWI/SNF complexes in adipogenesis, we first depleted endogenous SMARCA4, the catalytic ATPase subunit, using the auxin-inducible degron (AID) system^26^. Primary preadipocytes were isolated from newborn pups of *Smarca4*^AID/AID^ mice, which were generated using CRISPR-mediated insertion of an AID tag in the C-terminus of endogenous *Smarca4* alleles. Cells were immortalized and infected with a retroviral vector expressing the Myc-tagged auxin receptor Tir1. In the presence of auxin, Tir1 binds to the AID tag and induces proteasome-dependent degradation of endogenous SMARCA4 (Fig 2A). As shown in Fig. 2B, auxin induced a rapid Tir1-dependent depletion of endogenous SMARCA4 protein within 4h. Depletion of SMARCA4 prevented the appearance of lipid droplets, indicating that SMARCA4 is essential for adipogenesis in cell culture (Fig 2C).

**Figure 2.**
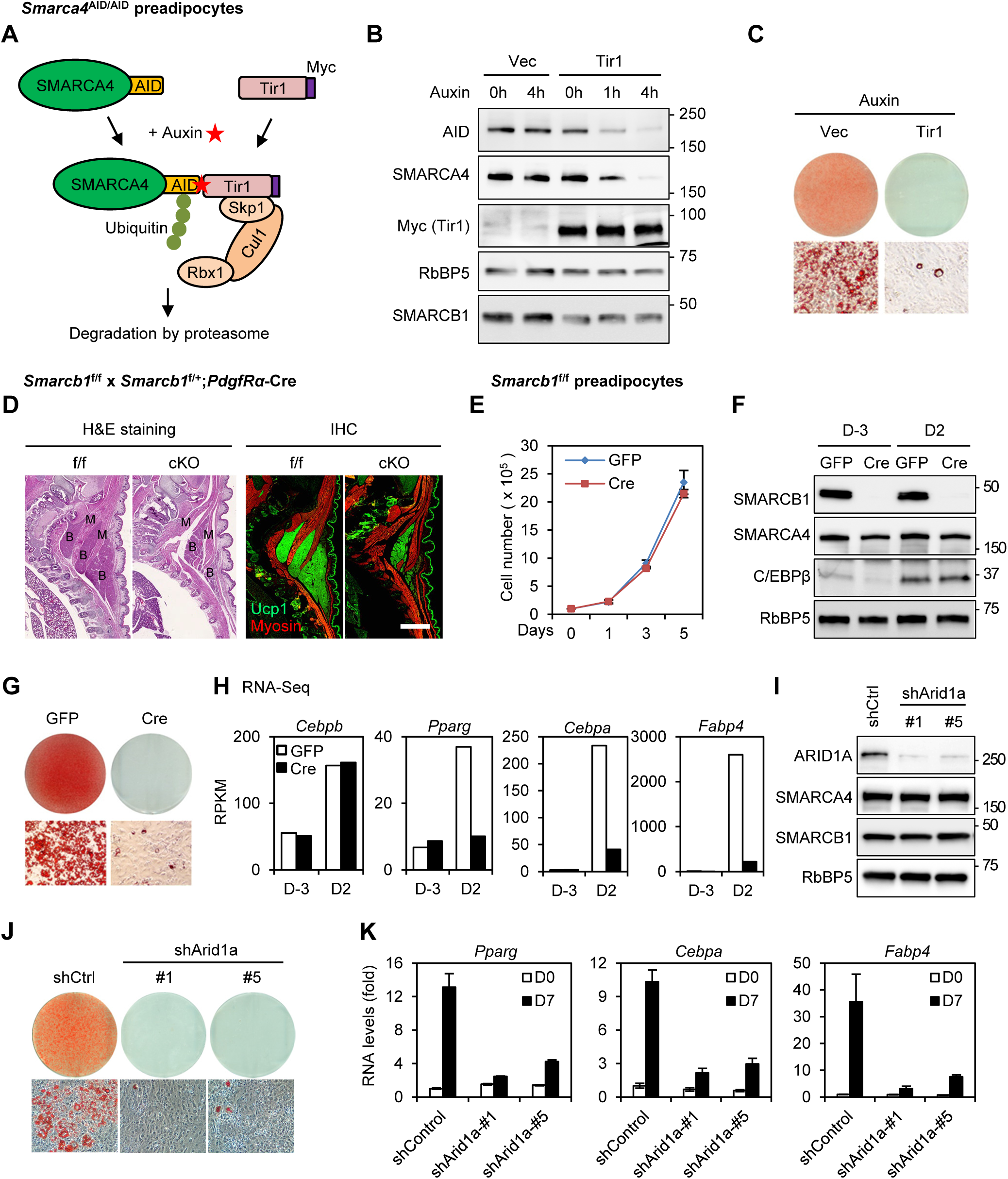
BAF subunits SMARCA4, SMARCB1 and ARID1A are required for adipogenesis. (**A-C**) Depletion of SMARCA4 by auxin-inducible degron (AID) system inhibits adipogenesis. **(A)** A schematic illustration of the auxin-induced depletion of endogenous SMARCA4 protein in Tir1-expressing *Smarca4*^AID/AID^ cells. TIR1, transport inhibitor response 1; Cul1, Cullin-1; Rbx1, RING-box protein 1; Skp1, S-phase kinase-associated protein 1. **(B)** Western blot analyses using antibodies indicated on the left. RbBP5 was used as a loading control. **(C)** Oil Red O staining at day 7 (D7) of adipogenesis. Cells were treated with auxin throughout the differentiation. **(D)** SMARCB1 is essential for adipogenesis *in vivo*. Sections of the interscapular area of E18.5 *Smarcb1*^f/f^ (f/f) and *Smarcb1*^f/f^;*PdgfRα-Cre* (cKO) embryos were stained with H&E (left panels) or with antibodies recognizing brown adipose tissue (BAT) marker Ucp1 (green) and skeletal muscle marker Myosin (red) (right panels). B, BAT; M, muscle. Scale bar = 1mm. (**E-H**) SMARCB1 is essential for adipogenesis in culture. SV40T-immortalized *Smarcb1*^f/f^ preadipocytes were infected with adenoviral GFP or Cre, followed by adipogenesis assays. **(E)** Deletion of *Smarcb1* does not affect cell growth rates. 1 x 10^5^ preadipocytes were plated and the cumulative cell numbers were determined for 5 days. **(F)** Before (D-3) and during (D2) adipogenesis, nuclear extracts were analyzed by Western blot using indicated antibodies. **(G)** Oil Red O staining at D7 of adipogenesis. **(H)** Expression of *Cebpb, Pparg, Cebpa* and *Fabp4* before (D-3) and during (D2) adipogenesis was determined using RNA-Seq. RPKM values indicate gene expression levels. (**I**-**K**) ARID1A is required for adipogenesis in culture. Preadipocytes were infected with lentiviral vectors expressing control (shCtrl) or *Arid1a* knockdown shRNA (shArid1a), followed by adipogenesis assays. **(I)** Western blot analysis of ARID1A in preadipocytes. **(J)** Oil Red O staining at D7 of adipogenesis. **(K)** qRT-PCR analysis of adipogenesis marker expression at indicated time points of adipogenesis.

Next, we examined the role of SMARCB1 in adipogenesis. We generated a conditional knockout (cKO) of SMARCB1 in preadipocytes by crossing *Smarcb1*^f/f^ mice with *PdgfRα-Cre* mice. The *PdgfRα-Cre* transgene is expressed in most adipocyte precursor cells^27^. *Smarcb1* cKO mice (*Smarcb1*^f/f^;*PdgfRα-Cre*) died shortly after birth, presumably due to severe defects in cranial development (Fig 2-S1A-D). Immunohistochemical analysis revealed that deletion of *Smarcb1* led to a severe reduction of interscapular brown adipose tissue (BAT) in E18.5 embryos (Fig 2D and Fig 2-S1E). Consistent with this phenotype, deletion of *Smarcb1* reduced the expression of adipogenesis markers *Pparg, Cebpa*, and *Fabp4* as well as BAT markers *Prdm16* and *Ucp1*, but not myogenesis markers (Fig 2-S1F). These findings indicate that SMARCB1 is required for adipose tissue development in mice.

To understand how SMARCB1 regulates adipogenesis, we isolated and immortalized *Smarcb1*^f/f^ preadipocytes. Deletion of *Smarcb1* in immortalized preadipocytes did not affect growth rates (Fig 2E). Consistent with previous data^9^, deletion of *Smarcb1* did not affect the expression of SMARCA4 nor the interaction between SMARCA4 and core subunit SMARCC2 (Fig 2F and Fig 2-S1G). However, deletion of *Smarcb1* blocked adipogenesis (Fig 2G). We performed RNA-Seq analysis before (D-3) and during (D2) adipogenesis. Using a 2.5-fold cut-off for differential expression, we defined 937 and 1,382 genes that were up- and down-regulated from D-3 to D2 of differentiation (Fig 2-S1H). Among the 937 up-regulated genes, 362 were induced in a SMARCB1-dependent manner. GO analysis showed that these genes were strongly associated functionally with fat cell differentiation and lipid metabolism (Fig 2-S1I). Deletion of *Smarcb1* did not affect the induction of pioneer TF C/EBPβ or the expression of other SWI/SNF subunits, but blocked the induction of adipocyte marker genes *Pparg, Cebpa*, and *Fabp4* (Fig 2F, 2H and Fig 2-S1J), suggesting that SMARCB1 is required for the induction of adipocyte genes downstream of C/EBPβ.

Because SMARCA4 and SMARCB1 are common subunits in BAF and PBAF, we wanted to clarify the function of each complex in adipogenesis. To determine the role of BAF in adipogenesis, we infected preadipocytes with lentiviral shRNA targeting BAF-specific subunit ARID1A. Knockdown of ARID1A did not affect SMARCA4 and SMARCB1 expression (Fig 2I), but it severely impaired adipogenesis (Fig 2J-K). To study the role of PBAF in adipogenesis, we generated a cKO of PBAF-specific subunit PBRM1 in precursor cells of brown adipocytes by crossing *Pbrm1*^f/f^ mice with *Myf5-Cre* mice. *Myf5-Cre* is specifically expressed in somitic precursor cells for both BAT and skeletal muscle in the interscapular region of mice^14^. *Pbrm1*^f/f^;*Myf5-Cre* (cKO) pups were born at the expected Mendelian ratio and survived to adulthood without obvious developmental defects (Fig 2-S2A-B). The adipose tissues isolated from cKO mice were similar in size and morphology to the control mice (Fig 2-S2C-D). Although the *Pbrm1* gene level was reduced by over 80% in the BAT of cKO mice, deletion of *Pbrm1* did not significantly affect the expression of adipogenesis markers *Pparg, Cebpa*, and *Fabp4* or BAT marker *Ucp1* (Fig 2-S2E). To further understand the role of PBRM1 in adipogenesis, we induced differentiation of primary brown preadipocytes isolated from cKO mice. Deletion of *Pbrm1* by Myf5-Cre had little effects on adipogenesis in culture (Fig 2-S2F-G). By RNA-Seq analysis at D7 of adipogenesis, we found that only a small number of genes were increased (0.4%) or decreased (0.4%) over 2.5-fold in *Pbrm1* cKO cells compared with control cells (Fig 2-S2H). Differentially regulated genes were not associated with adipocyte differentiation or lipid metabolism (Fig 2-S2I-J). Taken together, these findings suggest that BAF, but not PBAF, is essential for adipogenesis.

### BAF co-localizes with LDTFs and MLL4 on cell type-specific enhancers during adipogenesis

To assess the genomic binding of BAF, we first selected SMARCA4^+^ AEs (31,349) or promoters (8,871), then examined changes in genomic binding of BAF subunits SS18 and ARID1A before (D-3) and during (D2) adipogenesis. While genomic binding of the common ATPase subunit SMARCA4 was found on both AEs and promoters, genomic bindings of SS18 and ARID1A were mainly localized on AEs and increased from D-3 to D2 (Fig 3A). Motif analysis of the top 3,000 BAF-bound (SMARCA4^+^ SS18^+^) AEs showed enrichment of AP-1 family TF motifs at D-3 and motifs of adipogenic TFs C/EBPs and ATF4 at D2 (Fig 3B and Fig 3-S1A-C). These results indicate that BAF is enriched on cell type-specific enhancers during adipogenesis.

**Figure 3.**
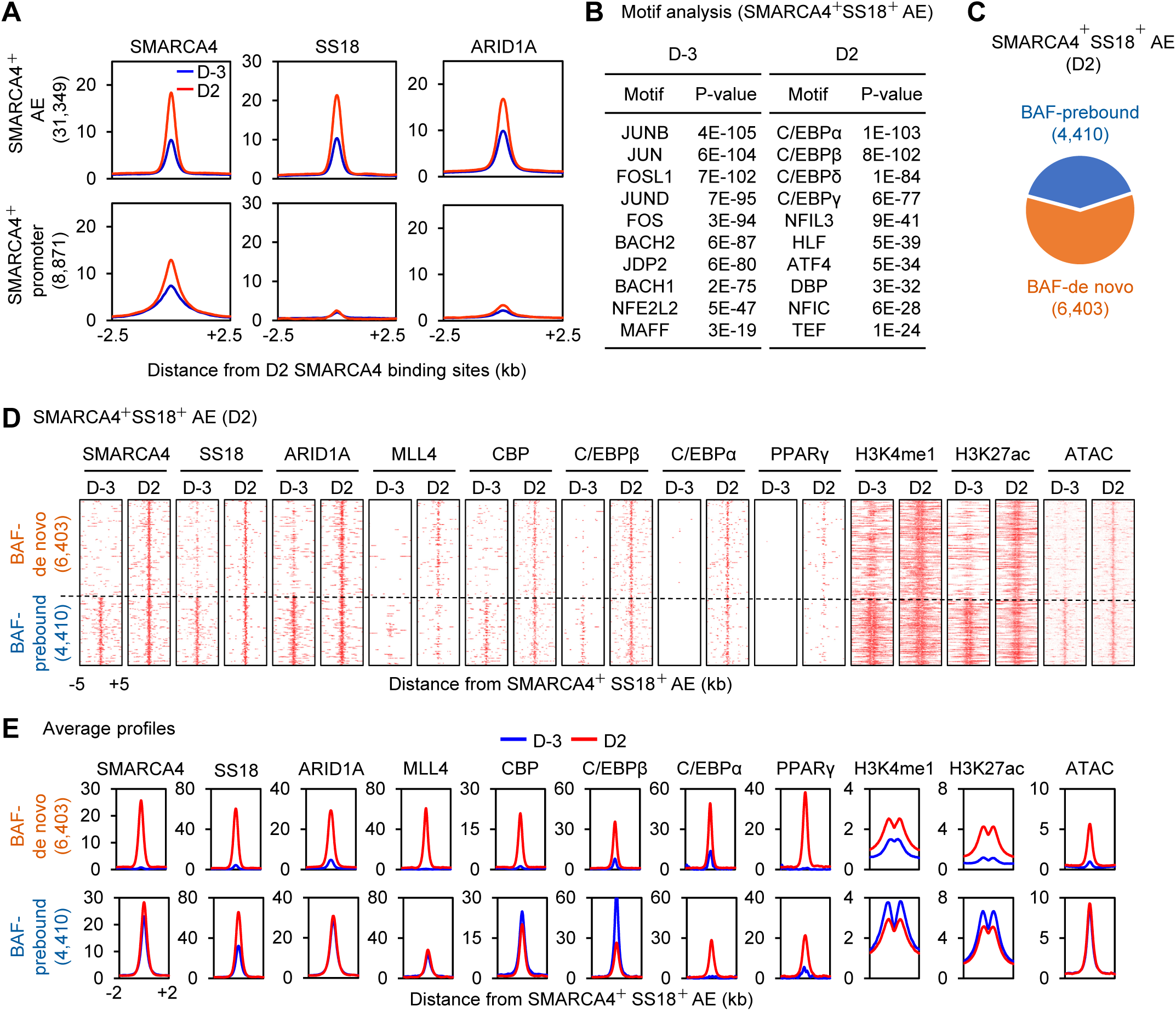
BAF co-localizes with LDTFs and MLL4 on active enhancers during adipogenesis. **(A)** Average binding profiles of SMARCA4, SS18, and ARID1A around the center of SMARCA4^+^ active enhancers (AEs) or promoters before (D-3) and during (D2) adipogenesis. Normalized read counts are shown. **(B)** Motif analysis of the top 3,000 SMARCA4^+^ SS18^+^ AEs. **(C)** Pie chart depicting BAF-binding (SMARCA4^+^ SS18^+^) AEs at D2 of adipogenesis. Among the 10,813 BAF-binding AEs at D2, 6,403 are *de novo* BAF-binding AEs (BAF-de novo), while 4,410 AEs are already bound by BAF (BAF-prebound) at D-3. (**D-E**) BAF subunits SMARCA4, SS18 and ARID1A colocalize with LDTFs (C/EBPβ, C/EBPα, and PPARγ), MLL4, CBP and open chromatin on AEs (H3K4me1^+^ H3K27ac^+^) at D2 of adipogenesis. The local epigenetic environment exhibits coordinated changes during differentiation. Heat maps (**D**) and average profiles (**E**) were aligned around the center of SMARCA4^+^ SS18^+^ AEs. Published ChIP-Seq data sets for MLL4, CBP, C/EBPβ, C/EBPα, and PPARγ were used (GSE74189)^18^.

To obtain insights into how LDTFs coordinate with BAF on AEs, we analyzed the genomic localization of adipogenic TFs C/EBPβ, C/EBPα and PPARγ, which are sequentially induced to activate adipocyte-specific genes in adipogenesis^28,29^. Among the 10,813 BAF-bound AEs during adipogenesis (D2), BAF was already bound to 4,410 of them in preadipocytes (D-3) and maintained the binding at D2 (BAF-prebound AEs). In contrast, 6,403 AEs exhibited emergent BAF binding at D2 but not at D-3 (BAF-de novo AEs) (Fig 3C). Consistent with motif analysis, BAF was highly co-localized with C/EBPβ, C/EBPα and PPARγ on AEs at D2 of adipogenesis. Particularly, C/EBPβ binding was detected on BAF-prebound AEs at D-3 but increased markedly from D-3 to D2 on BAF-de novo AEs (Fig 3D-E). Since MLL4 and CBP also colocalize with LDTFs on cell type-specific AEs during adipogenesis^18^, we assessed the co-occupancy among BAF, MLL4 and CBP. We found that MLL4 and CBP co-localized with BAF on both BAF-prebound and BAF-de novo AEs. While MLL4 and CBP binding on BAF-prebound AEs remained largely unchanged from D-3 to D2 of adipogenesis, their binding on BAF-de novo AEs was induced. Accordingly, the levels of MLL3/MLL4-catalyzed H3K4me1 and CBP/p300-catalyzed H3K27ac also increased on BAF-de novo AEs from D-3 to D2 (Fig 3D-E). Colocalization of BAF with LDTFs, MLL4 and CBP on BAF-de novo AEs was also found on *Pparg* and *Cebpa* gene loci during adipogenesis (Fig 3-S1D-E). ATAC-Seq analysis revealed that chromatin accessibility was highly correlated with BAF binding intensities on AEs during adipogenesis. The induction of chromatin accessibility on BAF-de novo AEs coincided with the increase in the binding of LDTFs, MLL4 and CBP on AEs (Fig 3D-E and Fig 3-S1D-E). These results indicate that BAF colocalizes with LDTFs, MLL4 and CBP on cell type-specific enhancers and promotes chromatin accessibility during differentiation.

At D2 of adipogenesis, about 40% (4,362/10,813) of BAF-bound AEs were occupied by MLL4 (Fig 3-S2A). Differential motif analysis of BAF-bound AEs revealed that motifs of LDTFs PPARγ and C/EBPs were enriched on MLL4^+^ AEs while motifs of the AP-1 family TFs were enriched on MLL4^-^ AEs at D2 (Fig 3-S2B). Heat maps and average profiles also showed that from D-3 to D2 of adipogenesis, the binding of LDTFs and BAF subunits (SMARCA4, SS18 and ARID1A) was more significantly increased on MLL4^+^ than MLL4^-^ AEs. Particularly, levels of CBP and H3K27ac were only induced on MLL4^+^ AEs (Fig 3-S2C and D). D2 MLL4^-^ AEs were already accessible before adipogenesis (D-3). In contrast, chromatin accessibility on D2 MLL4^+^ AEs was dramatically increased from D-3 to D2 (Fig 3-S2C and D). These results suggest the involvement of MLL4 in enhancer activation by BAF.

To test the possibility of physical interaction between BAF and MLL4, we identified proteins associated with endogenous MLL4 and UTX, a subunit of the MLL4 complex, in mouse embryonic stem cells (ESCs) by mass spectrometry. Consistent with previous results^30^, while immunoprecipitation with an anti-UTX antibody pulled down both MLL3 and MLL4 as well as other components of MLL3/MLL4 complexes, MLL3 was not co-immunoprecipitated by an anti-MLL4 antibody. Notably, several SWI/SNF subunits, including SMARCA4, SMARCC1, SMARCD1, SMARCB1 and SMARCE1, were also pulled down by anti-MLL4 and/or anti-UTX antibody from ESC nuclear extracts (Fig 3-S3A). We also confirmed that endogenous SMARCA4 and SMARCB1 are associated with endogenous MLL4 and UTX, but not the MLL1 complex subunit Menin, in HEK293T cells and preadipocytes at D2 of adipogenesis (Fig 3-S3B-C). Together, these findings suggest that BAF cooperates with MLL4 to activate cell type-specific enhancers during differentiation.

### Reciprocal regulation between BAF and MLL4 on active enhancers during adipogenesis

To study the relationship between BAF and MLL4 on cell type-specific enhancers, we assessed their genomic binding on adipogenic enhancers at D2 of differentiation. Adipogenic enhancers were defined as AEs that are bound by C/EBPs or PPARγ (Fig 4A). This set of enhancers are mostly de novo. We first asked whether MLL4 is required for BAF recruitment to adipogenic enhancers during differentiation. To eliminate the compensatory effect of MLL3, we deleted *Mll4* in *Mll3*^-/-^*Mll4*^f/f^ cells^14^. ChIP-Seq revealed that deletion of *Mll4* prevented genomic binding of SMARCA4 and SS18 on adipogenic enhancers. Consistently, ATAC-Seq showed that chromatin accessibility at these enhancers was also significantly reduced in *Mll4* KO cells (Fig 4A-B). Reduced binding of BAF on adipogenic enhancers in *Mll4* KO cells was confirmed on *Pparg* and *Fabp4* gene loci (Fig 4-S1A-B).

**Figure 4.**
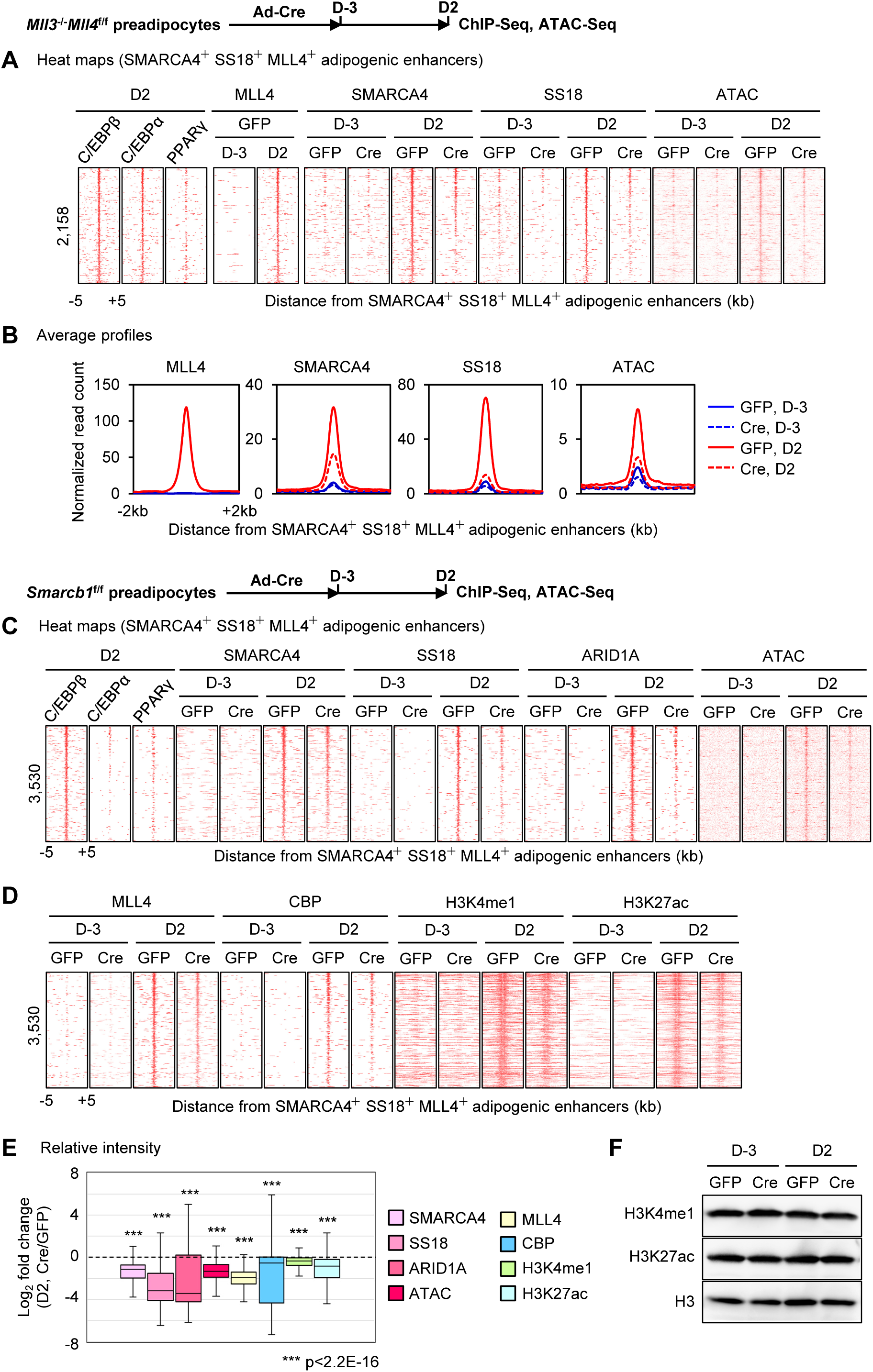
Reciprocal regulation between BAF and MLL4 on active enhancers during adipogenesis. *Mll3*^-/-^*Mll4*^f/f^ or *Smarcb1*^f/f^ preadipocytes were infected with adenoviruses expressing GFP or Cre, followed by adipogenesis assays. Cells were collected before (D-3) and during (D2) adipogenesis for ChIP-Seq, ATAC-Seq and Western blot analyses. (**A-B**) Decreased SMARCA4 and SS18 binding and chromatin accessibility on adipogenic enhancers in *Mll4* KO (Cre) cells during adipogenesis. Adipogenic enhancers were defined as active enhancers bound by either C/EBPβ, C/EBPα, or PPARγ at D2 of adipogenesis. Published ChIP-Seq data sets for C/EBPβ, C/EBPα, and PPARγ were used (GSE74189) ^18^. Heat maps (**A**) and average profiles (**B**) are shown around SMARCA4^+^ SS18^+^ MLL4^+^ adipogenic enhancers. Adipogenic enhancers shown in the heat maps were ranked by the binding intensity of SMARCA4 at the center in D2 control (GFP) cells. (**C-F**) Deletion of *Smarcb1* reduces binding of BAF subunits (SMARCA4, SS18, and ARID1A) as well as chromatin opening, and impairs MLL4 binding and activation of adipogenic enhancers. Adipogenic enhancers at D2 of adipogenesis were re-defined using ChIP-Seq data sets obtained in *Smarcb1*^f/f^ cells. **(C)** Heat maps for ChIP-Seq of BAF subunits (SMARCA4, SS18, and ARID1A) and ATAC-Seq around the center of SMARCA4^+^ SS18^+^ MLL4^+^ adipogenic enhancers. **(D)** Heat maps for ChIP-Seq of MLL4, CBP, H3K4me1 and H3K27ac on adipogenic enhancers. Adipogenic enhancers shown in the heat maps were ranked by the intensity of SMARCA4 at the center in D2 control (GFP) cells. **(E)** Fold changes of intensities between control (GFP) and *Smarcb1* KO (Cre) cells on adipogenic enhancers are shown in box plots. Statistical significance levels are indicated (Wilcoxon signed rank test, one-sided). **(F)** Western blot analyses of H3K4me1 and H3K27ac in control and *Smarcb1* KO cells during adipogenesis. Total histone H3 was used as a loading control.

We next evaluated whether BAF contributes to the genomic binding of MLL4 on adipogenic enhancers during differentiation. ChIP-Seq and ATAC-Seq revealed that *Smarcb1* deletion in preadipocytes reduced genomic binding of SMARCA4, SS18 and ARID1A, as well as chromatin accessibility, on adipogenic enhancers at D2 of adipogenesis (Fig 4C and 4E). Moreover, deletion of *Smarcb1* reduced binding of MLL4 and CBP on adipogenic enhancers, indicating that BAF is required for the activation of cell type-specific enhancers (Fig 4D-E). H3K4me1 and H3K27ac levels also decreased on adipogenic enhancers in *Smarcb1* KO cells, although global levels of these enhancer marks were unchanged (Fig 4F). Reduced binding of MLL4 on adipogenic enhancers in *Smarcb1* KO cells was confirmed on the *Pparg* gene locus (Fig 4-S1C). Together, these findings reveal reciprocal regulation between BAF and MLL4 on adipogenic enhancers and suggest that BAF and MLL4 coordinate to activate cell type-specific enhancers during differentiation.

### BAF and MLL4 reciprocally promote each other’s binding to maintain active enhancers in preadipocytes

Because we observed interdependent genomic binding of BAF and MLL4 during adipogenesis, we next wondered whether such a relationship is also present in undifferentiated preadipocytes. To examine the genomic association of BAF and MLL4 on AEs in undifferentiated cells, we identified 6,304 BAF^+^ MLL4^+^ AEs that were shared by *Mll3*^-/-^*Mll4*^f/f^ and *Smarcb1*^f/f^ preadipocytes. Deletion of *Mll4* in *Mll3*^-/-^ *Mll4*^f/f^ preadipocytes reduced ARID1A binding and H3K27ac levels, but not SMARCA4 binding, on BAF^+^ MLL4^+^ AEs in preadipocytes (Fig 5A-B). Next, we investigated whether BAF is required for maintaining MLL4 binding on AEs in preadipocytes. While deletion of *Smarcb1* had little effect on maintaining genomic binding of SMARCA4 or chromatin accessibility, it reduced MLL4 binding on BAF^+^ MLL4^+^ AEs (Fig 5C-D). Decreased MLL4 binding on BAF^+^ MLL4^+^ AEs was also observed in SMARCA4-depleted preadipocytes in addition to widespread reduction in chromatin accessibility (Fig 5E-F). Together, these data suggest a reciprocal regulation between BAF and MLL4 in promoting their bindings on AEs in undifferentiated cells.

**Figure 5.**
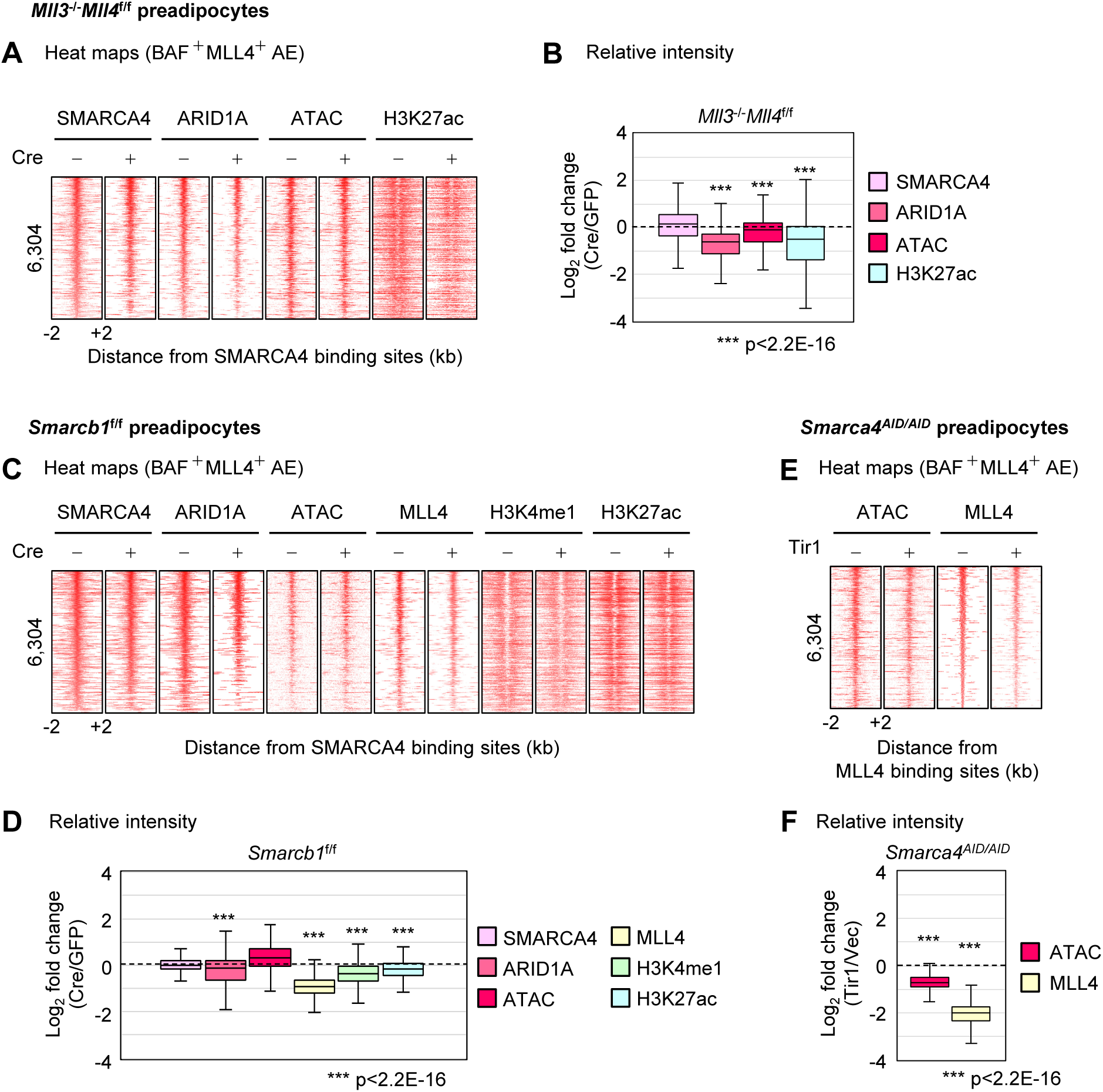
BAF and MLL4 reciprocally promote each other’s binding to maintain active enhancers in preadipocytes. *Mll3*^-/-^*Mll4*^f/f^ or *Smarcb1*^f/f^ preadipocytes were infected with adenoviruses expressing GFP or Cre. Sub-confluent cells were collected for ChIP-Seq and ATAC-Seq. BAF^+^ MLL4^+^ active enhancers (AEs) were defined as those bound by SMARCA4, ARID1A and MLL4. (**A-B**) Deletion of *Mll4* reduces ARID1A binding on BAF^+^ MLL4^+^ AEs in preadipocytes. **(A)** AEs shown in the heat maps were ranked by the intensity of SMARCA4 at the center in control (GFP) cells. **(B)** Fold changes of intensities between control and *Mll4* KO (Cre) cells on AEs in preadipocytes are shown in box plots. Statistical significance levels are indicated (Wilcoxon signed rank test, one-sided). (**C-D**) SMARCB1 is required for maintaining MLL4 binding on BAF^+^ MLL4^+^ AEs in preadipocytes. **(C)** AEs shown in the heat maps were ranked by the intensity of SMARCA4 at the center in control cells. **(D)** Fold changes of intensities between control (GFP) and *Smarcb1* KO (Cre) cells on AEs in preadipocytes are shown in box plots. Statistical significance levels are indicated (Wilcoxon signed rank test, one-sided). (**E-F**) SMARCA4 is required for maintaining MLL4 binding on BAF^+^ MLL4^+^ AEs in preadipocytes. *Smarca4*^AID/AID^ preadipocytes were infected with retroviruses expressing Tir1 or vector (Vec) only, followed by auxin (0.5 mM) treatment for 4h. Cells were then collected for ATAC-Seq and ChIP-Seq of MLL4. **(E)** AEs shown in the heat maps were ranked by the intensity of MLL4 at the center in control cells. **(F)** Fold changes of intensities between SMARCA4-depleted (Tir1) and control (Vec) cells on AEs in preadipocytes are shown in box plots. Statistical significance levels are indicated (Wilcoxon signed rank test, one-sided).

### MLL4 is required for BAF binding on C/EBPβ-activated enhancers

Interdependent genomic binding of BAF and MLL4 on AEs during adipogenesis could be due to indirect consequences of differentiation defect and failed induction of LDTFs. To evaluate the primary effect of MLL4 on LDTF-activated enhancers, we used *Mll3*^-/-^*Mll4*^f/f^ preadipocytes expressing ectopic C/EBPβ, which is a pioneer factor that activates a subset of adipogenic enhancers in preadipocytes without inducing differentiation^14,25^. Neither ectopic expression of C/EBPβ nor deletion of *Mll4* affected expression levels of BAF subunits SMARCA4, SS18 and ARID1A in preadipocytes (Fig 6A). We identified 571 C/EBPβ-activated enhancers that were bound by MLL4 and BAF subunits SMARCA4 and ARID1A (see Materials and Methods). Among these enhancers, 324 displayed ectopic C/EBPβ-induced *de novo* BAF binding (BAF-de novo), whereas the remaining 247 were pre-marked by BAF in control cells before ectopic expression of C/EBPβ (BAF-prebound) (Fig 6B). As expected, motifs of C/EBPs were enriched on both BAF-de novo and BAF-prebound C/EBPβ-activated enhancers (Fig 6C). Consistent with our previous finding, deletion of *Mll4* markedly decreased H3K27ac levels on C/EBPβ-activated enhancers. Importantly, deletion of *Mll4* not only prevented C/EBPβ-induced *de novo* BAF binding (as measured by SMARCA4 and ARID1A), but also compromised genomic binding of SMARCA4 and ARID1A on BAF-prebound enhancers, despite little changes in C/EBPβ binding (Fig 6D and E). These results indicate that MLL4 is required for genomic binding of BAF on C/EBPβ-activated enhancers and suggest a direct role of MLL4 in mediating BAF localization on enhancers.

**Figure 6.**
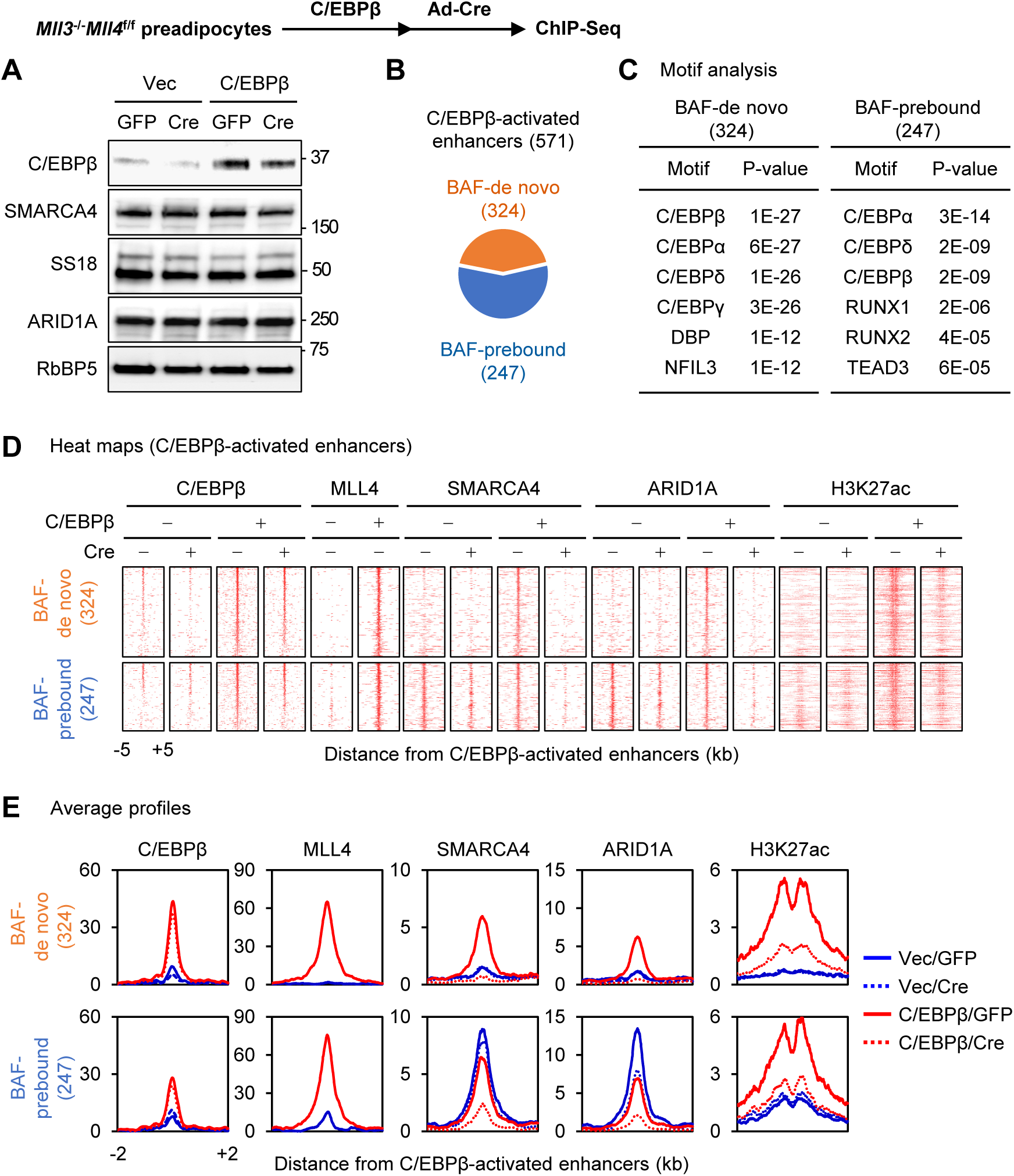
MLL4 is required for BAF binding on C/EBPβ-activated enhancers. *Mll3*^-/-^*Mll4*^f/f^ preadipocytes were infected with retroviruses expressing C/EBPβ or vector (Vec), and then infected with adenoviruses expressing GFP or Cre. Sub-confluent cells were collected without inducing adipogenesis for Western blot and ChIP-Seq analyses. Published ChIP-Seq data sets for C/EBPβ, MLL4, H3K27ac were used (GSE50466)^14^. **(A)** Western blot analyses of BAF subunits (SMARCA4, SS18, and ARID1A) and C/EBPβ expression in control (GFP) and *Mll4* KO (Cre) preadipocytes. **(B)** Among the 571 C/EBPβ-activated enhancers in preadipocytes, BAF (SMARCA4 and ARID1A) binding was induced on 324 AEs only after ectopic expression of C/EBPβ (BAF-de novo), while 247 AEs were already bound by BAF before ectopic expression of C/EBPβ in preadipocytes (BAF-prebound). **(C)** Motif analysis of BAF-de novo and BAF-prebound AEs. (**D-E**) Deletion of *Mll4* does not affect C/EBPβ binding but reduces BAF binding on C/EBPβ-activated enhancers. Heat maps (**D**) and average profiles (**E**) around C/EBPβ-activated enhancers are shown. Enhancers shown in the heat maps were ranked by the intensity of C/EBPβ at the center in control cells expressing ectopic C/EBPβ.

### BAF regulates MLL4 binding on C/EBPβ-activated enhancers

We next asked whether BAF is required for MLL4 binding on C/EBPβ-activated enhancers. For this purpose, we infected *Smarca4*^AID/AID^ preadipocytes with retroviruses expressing C/EBPβ followed by retroviruses expressing Tir1. Cells were then treated with auxin for 4h to deplete endogenous SMARCA4 protein. Western blot showed that acute depletion of SMARCA4 did not affect the expression levels of ectopic C/EBPβ in preadipocytes (Fig 7A). Among 449 C/EBPβ-activated enhancers that we identified, C/EBPβ binding was unchanged at 220 AEs (SMARCA4-independent), while C/EBPβ occupancy was reduced at the remaining 229 AEs (SMARCA4-dependent) upon SMARCA4 depletion (Fig 7B). Only SMARCA4-independent C/EBPβ binding AEs were enriched with C/EBP motifs while SMARCA4-dependent ones were enriched with motifs of AP1 family TFs, suggesting that SMARCA4-independent C/EBPβ binding on AEs is direct while SMARCA4-dependent C/EBPβ binding is through tethering to AP1-bound AEs (Fig 7C). To assess the direct effect of SMARCA4 on MLL4 recruitment to C/EBPβ-activated enhancers, we focused on SMARCA4-independent AEs. Despite intact C/EBPβ binding, SMARCA4 depletion led to decreased MLL4 binding and chromatin accessibility on these AEs (Fig 7D-E). These data demonstrate that SMARCA4 is required for MLL4 binding on C/EBPβ-activated enhancers. Together, our data suggest an interdependent relationship between BAF and MLL4 in pioneer LDTF-mediated enhancer activation (Fig 7F).

**Figure 7.**
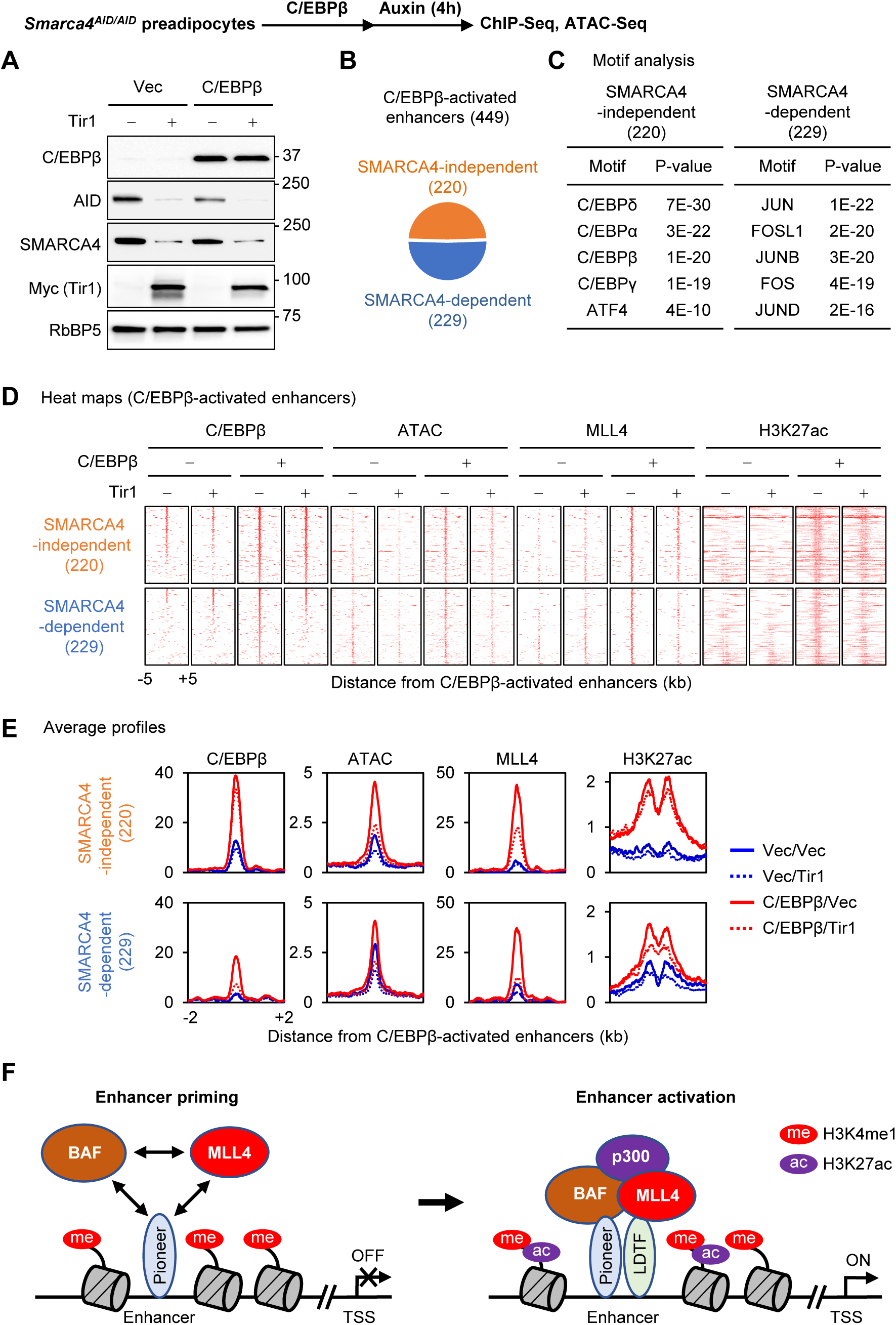
BAF regulates MLL4 binding on C/EBPβ-activated enhancers. *Smarca4*^AID/AID^ preadipocytes were infected with retroviruses expressing C/EBPβ or vector (Vec), and then infected with retroviruses expressing Tir1. Sub-confluent cells were treated with auxin for 4h and collected for Western blot, ChIP-Seq and ATAC-Seq analyses. **(A)** Western blot analyses using antibodies indicated on the left. **(B)** Among the 449 C/EBPβ-activated enhancers in preadipocytes, 220 exhibited SMARCA4-independent C/EBPβ binding, while the remaining 229 showed SMARCA4-dependent C/EBPβ binding. **(C)** Motif analysis of SMARCA4-independent and -dependent C/EBPβ-activated enhancers in preadipocytes. (**D-E**) Auxin-induced, Tir1-dependent acute depletion of SMARCA4 reduces MLL4 binding on C/EBPβ-activated enhancers in preadipocytes. Heat maps (**D**) and average profiles (**E**) around C/EBPβ-activated enhancers are shown. Enhancers shown in the heat maps were ranked by the intensity of C/EBPβ at the center in control cells expressing ectopic C/EBPβ. (**F**) A model depicting the interdependent relationship between BAF and MLL4 in promoting pioneer TF- and LDTF-dependent cell type-specific enhancer activation.

## Discussion

Using adipogenesis as a model system, we provide several lines of evidence to support that BAF cooperates with MLL4 to promote LDTF-dependent enhancer activation and cell type-specific gene expression during cell differentiation. We found that BAF, similar to MLL4, is required for adipogenesis by using conditional knockout mice and derived cells. BAF, but not PBAF, colocalizes with LDTFs and MLL4 on cell type-specific enhancers. Similar to MLL4, BAF is essential for enhancer activation and cell type-specific gene induction during adipogenesis. By depleting BAF subunits SMARCA4 and SMARCB1 as well as MLL4 in cells, we demonstrated that BAF and MLL4 reciprocally regulate one another’s binding on AEs before and during adipogenesis. Finally, by evaluating pioneer factor C/EBPβ-activated enhancers without inducing differentiation, we showed direct evidence for an interdependent relationship between BAF and MLL4 on activating cell type-specific enhancers.

### Function of SWI/SNF complexes in adipogenesis

Several SWI/SNF subunits have been shown to regulate adipogenesis in culture^28^. Ectopic expression of dominant-negative SMARCA4 interferes with PPARγ-, C/EBPα- and C/EBPβ-stimulated adipogenesis in fibroblasts^24^. Knockdown of *Smarcb1* in 3T3-L1 preadipocytes inhibits adipogenesis^31^. Using the AID system, we demonstrated that endogenous SMARCA4 is essential for adipogenesis. We showed the essential role of SMARCB1 in adipogenesis of SV40T-immortalized preadipocytes. Immortalization by SV40T, which inactivates tumor suppressors p53 and Rb^32^, prevents growth defects caused by *Smarcb1* deletion^33,34^, and enables the study of SMARCB1 in preadipocyte differentiation. Further, we showed that SMARCB1 is required for adipose tissue development in mice. We also observed that the BAF-specific subunit ARID1A is required for adipogenesis in preadipocytes. Together, these findings demonstrate a critical role of BAF in adipogenesis. Our observations that the PBAF-specific PBRM1 is dispensable for adipogenesis *in vitro* and *in vivo* provide an example of distinct functions of BAF and PBAF in regulating cell differentiation and development. The role of GBAF in adipogenesis remains to be determined.

### Distinct genomic localization of BAF, PBAF and GBAF

Using adipogenesis as a model system for cell differentiation, we confirmed previous observations on the distinct genomic localization of BAF, PBAF and GBAF in ESCs and in cancer cells: BAF to AEs, PBAF to promoters and GBAF to CTCF binding sites^11,12,35^. Although SMARCA4 and SS18 were reported to physically associate with both BAF and GBAF complexes, we found that on AEs, the vast majority of them are associated with BAF. We also showed that BAF is the major SWI/SNF complex on AEs and that BAF binding sites dynamically change during adipogenesis. Motifs of AP-1 family TFs are enriched on BAF-bound AEs in preadipocytes, consistent with a recent finding that AP-1 can recruit BAF to establish accessible chromatin in fibroblasts^36^. Although GBAF is enriched on CTCF binding sites, it also colocalizes with BAF on a subset of AEs during adipogenesis. The role of GBAF on AEs in cell differentiation remains unclear.

### Reciprocal regulation between BAF and MLL4 in enhancer activation

We identified 4,410 BAF-prebound AEs and 6,403 BAF-de novo AEs at D2 of adipogenesis (Fig 3D-E). While BAF-prebound enhancers were constitutively active, BAF-de novo enhancers were activated from D-3 (preadipocytes) to D2 of adipogenesis. BAF-de novo AEs were more enriched with MLL4 binding (63.4%; 4,065/6,403), compared with BAF-prebound ones (6.7%; 297/4,410) at D2 of adipogenesis. In contrast to MLL4^-^ AEs, MLL4^+^ ones showed markedly increased binding of BAF and LDTFs, chromatin accessibility and enhancer activation during adipogenesis (Fig 3-S2). These observations suggest that BAF cooperates with MLL4 to activate cell type-specific enhancers during cell fate transition.

It has been reported that MLL3/MLL4-catalyzed H3K4me1 can directly recruit BAF complex to enhancers and that knockout of MLL3/MLL4 in ESCs reduces SMARCA4 binding on enhancers^37^, which imply that MLL3/MLL4 are necessary for BAF recruitment to enhancers. On the other hand, a more recent study showed that BAF does not require pre-marked H3K4me1 for binding on enhancers and that ectopic expression of SMARCA4 in SMARCA4/SMARCA2-deficient cancer cells establishes MLL3/MLL4 binding on AEs bound by BAF^38^, suggesting that BAF plays a critical role in MLL3/MLL4 binding on enhancers. Through the use of several gene knockout and protein depletion systems, we establish a bidirectional relationship between BAF and MLL4 on AEs in preadipocytes and during adipogenesis, which rectifies seemingly conflicting results from previous separate studies. Reciprocal regulation between BAF and MLL4 is also supported by their physical association observed in this study and reported in the literature^39^. A recent study further showed that UTX, a component of MLL3/MLL4 complex, physically and functionally associates with SMARCA4 to maintain chromatin accessibility in myeloid cells^40^. Whether BAF and MLL4 regulate each other on AEs through physical interaction of their components or enhancer histone marks remains to be determined.

### Interplay of BAF and MLL4 promotes pioneer factor-dependent enhancer activation

Pioneer factors such as OCT4 and C/EBPβ physically associate with SMARCA4 and other BAF subunits in cells^23,41^. Pioneer factors recruit SMARCA4 to induce chromatin accessibility to facilitate enhancer activation^42^. We reported previously that OCT4 and C/EBPβ physically associate with the MLL4 complex in cells and that ectopic C/EBPβ recruits MLL4 to activate a subset of C/EBPβ binding adipogenic enhancers in undifferentiated cells^14,20^. It is worth noting that MLL4 loss does not affect C/EBPβ binding on AEs. In contrast, acute depletion of SMARCA4 reduces C/EBPβ occupancy on AEs, consistent with recent reports on the role of BAF in the binding of pioneer factors on enhancers^42,43^. Our findings in the current study further suggest a model for reciprocal regulation between BAF and MLL4 on C/EBPβ-activated enhancers. The pioneer factor C/EBPβ recruits BAF to remodel chromatin to enable MLL4 binding on enhancers. Meanwhile, C/EBPβ recruits MLL4 to facilitate BAF binding on enhancers. BAF and MLL4 may cooperate with C/EBPβ to maintain chromatin accessibility and active enhancer landscape and facilitate other LDTFs binding (Fig 7F). Thus, BAF and MLL4 form a positive feedback loop to activate enhancers and lineage-specific gene expression during cell differentiation.

## Materials and Methods

### Plasmids and antibodies

The retroviral plasmids pBABEneo-SV40LT and pWZLhygro-C/EBPβ have been described^14^. The homemade antibodies anti-UTX^44^ and anti-MLL4^45^ have been described. Anti-SMARCB1/SNF5/INI1 (A-5, sc-166165), anti-SMARCC2/BAF170 (E-6, sc-17838X), anti-ARID2/BAF200 (E-3, sc-166117X), anti-C/EBPβ (C-19, sc-150X), anti-C/EBPα (144AA, sc-61X) and anti-PPARγ (H-100, sc-7196X) were from Santa Cruz. Anti-RbBP5 (A300-109A) was from Bethyl Laboratories. Anti-SMARCA4/BRG1 (ab110641) and anti-H3K27ac (ab4729) were from Abcam. Anti-ARID1A/BAF250A (D2A8U, #12354), anti-SS18 (D6I4Z, #21792) and anti-CBP (D6C5, #7389) were from Cell Signaling. Anti-BRD9 (#61537) was from Active Motif. Anti-H3K4me1 (13-0040) was from EpiCypher.

### Mouse experiments

To generate *Smarcb1*^f/f^;*PdgfRα-Cre* mice, *Smarcb1*^f/f^ mice^46^ obtained from Charles W.M. Roberts (St. Jude Children’s Research Hospital, Memphis, TN) were crossed with *PdgfRα-Cre* mice (Jackson No.013148). *Pbrm1*^f/f^ mice (Jackson No. 029049) were crossed with *Myf5-Cre* mice (Jackson No. 007893) to get *Pbrm1*^f/f^;*Myf5-Cre* mice. Generation of *Smarca4*^AID/AID^ mice will be described in a separate manuscript. Histology and immunohistochemistry analyses of E18.5 embryos were done as described^14^. Anti-Myosin (1:20 dilution, MF20; Developmental Studies Hybridoma Bank) and anti-UCP1 (1:400 dilution, ab10983; Abcam) were used. For fluorescent secondary antibodies, anti-mouse Alexa Fluor 488 and anti-rabbit Alexa Fluor 555 (Life Technologies, Carlsbad, CA, USA) were used. All mouse work was approved by the Animal Care and Use Committee of NIDDK, NIH.

### Isolation and immortalization of primary preadipocytes, virus infection and adipogenesis assay

Primary preadipocytes were isolated from interscapular BAT of newborn pups and were immortalized with retroviral vectors expressing SV40T as described^45^. Adenoviral infection of preadipocytes was done at 50 moi. Cells were routinely cultured in DMEM plus 10% FBS. For adipogenesis assays, preadipocytes were plated at a density of 1 × 10^5^ in each well of 6-well plates in growth medium (DMEM plus 10% FBS) 4 days before induction of adipogenesis. At day 0, cells were fully confluent and were treated with differentiation medium (DMEM plus 10% FBS, 0.1 μM insulin and 1 nM T3) supplemented with 0.5 mM IBMX, 1 μM DEX, and 0.125 mM indomethacin. Two days later, cells were changed to the differentiation medium. Cells were replenished with fresh medium at 2-day intervals. Fully differentiated cells were either stained with Oil Red O or subjected to gene expression analyses.

### The auxin-inducible degron (AID) system for SMARCA4 depletion

*Smarca4*^AID/AID^ preadipocytes were isolated from interscapular BAT of *Smarca4*^AID/AID^ newborn pups and immortalized with SV40T. Immortalized *Smarca4*^AID/AID^ preadipocytes were infected with retroviral vector expressing the auxin receptor Tir1 of rice (pBabePuro-OsTir1-9Myc, Addgene #80074). To induce degradation of SMARCA4, auxin (indole-3-acetic acid sodium salt, Sigma-Aldrich #I5148) was added to the culture medium in a final concentration of 0.5 mM for indicated times.

### shRNA knockdown

For shRNA transduction, individual shRNAs targeting mouse *Arid1a* (TRCN0000238303, shArid1a-#1; TRCN0000238306, shArid1a-#5) were obtained from Sigma Mission shRNA library (Sigma-Aldrich). Immortalized *Smarcb1*^f/f^ preadipocytes were infected with lentiviral shRNA targeting *Arid1a* or control virus alone for 24h. Infected cells were selected with puromycin (2 μg/mL) for 4 days before performing further experiments.

### Western blot and immunoprecipitation

Nuclear proteins were extracted using the nuclear extract preparation method as described previously^19^. Briefly, cells were washed with cold PBS, resuspended in buffer A (10 mM HEPES, pH 7.9, 1.5 mM MgCl_2_, 10 mM KCl and 0.1% NP40) supplemented with protease inhibitors (Roche), 0.5 mM DTT and 0.2 mM phenylmethylsulfonyl fluoride (PMSF), and incubated on ice for 10 min. After centrifugation at 1,000g, nuclei were resuspended in buffer C (20 mM HEPES, pH 7.9, 1.5 mM MgCl_2_, 420 mM NaCl, 0.2 mM EDTA and 25% glycerol) supplemented with protease inhibitors, 0.5 mM DTT and 0.2 mM PMSF. Nuclear extracts were separated using 4–15% Tris-Glycine gradient gels (Bio-Rad Laboratories). For Western blotting of MLL4, nuclear extracts were separated using 3–8% Tris-Acetate gradient gels (Invitrogen). Total proteins on the gel were transferred to a PVDF membrane (Bio-Rad Laboratories). The membranes were probed using specific antibodies.

For co-immunoprecipitation (IP), nuclear proteins were extracted from HEK293T and preadipocytes, diluted with an IP Buffer (50 mM Tris-HCl, pH 7.5, 1 mM EDTA, 150 mM NaCl, and 1% Triton X-100) supplemented with protease inhibitors, 1 mM DTT and 1 mM PMSF, and incubated with anti-MLL4, anti-UTX, anti-SMARCA4 or anti-SMARCB1 antibodies overnight at 4 °C. Samples were additionally incubated with Dynabeads Protein A (Life Technologies, 10008D) at 4 °C for 2 h. Beads were washed three times with an IP buffer and the immunoprecipitates were eluted with 20 μLof 1x sample buffer (NuPage LDS buffer, Thermo Scientific) including 100 mM DTT.

### RNA isolation and qRT-PCR analysis

Total RNA was extracted using TRIzol (Life Technologies) and reverse transcribed using ProtoScript II first-strand cDNA synthesis kit (NEB, E6560), following manufacturer’s instructions. Quantitative RT-PCR (qRT-PCR) was performed in duplicate with the Luna^®^ Universal qPCR Master Mix (NEB, M3003) using QuantStudio 5 Real-Time PCR System (Thermo Fisher). PCR amplification parameters were 95 °C (3 min) and 40 cycles of 95 °C (15 s), 60 °C (60 s), and 72 °C (30 s). Primer sequences are available upon request. Statistical significance was calculated using the two-tailed unpaired *t*-test on two experimental conditions.

### RNA-Seq library preparation

Total RNA was subjected to mRNA purification using the NEBNext Poly(A) mRNA Magnetic Isolation Module (NEB, E7490). Isolated mRNAs were reverse-transcribed into double stranded cDNA and subjected to sequencing library construction using the NEBNext Ultra™ II RNA Library Prep Kit for Illumina (NEB, E7770) according to the manufacturer’s instructions. RNA libraries were sequenced on Illumina HiSeq 3000.

### ChIP and ChIP-Seq library preparation

Chromatin immunoprecipitation (ChIP) and ChIP-Seq were done as described^47^. Briefly, cells were cross-linked with 1.5% formaldehyde for 10 min and quenched by 125 mM glycine for 10 min. 10 million fixed cells were resuspended in ice cold buffer containing 5mM PIPES, pH 7.5, 85mM KCl, 1% NP-40 and protease inhibitors, incubated on ice for 15 min, and centrifuged at 500 g for 5 min at 4°C. Nuclei were resuspended with 1mL buffer containing 50mM Tris-HCl, pH8.0, 10mM EDTA, 0.1% SDS and protease inhibitors, and subjected to sonication. Sheared chromatin was clarified by centrifugation at 13,000 g for 10 min at 4°C. The supernatant was transferred to a new tube and further supplemented with 1% Triton-X100, 0.1% sodium deoxycholate and protease inhibitors. 2% of the mixture was set aside as input and 20ng of spike-in chromatin (Active Motif, #53083) was added to the rest. For each ChIP, 4 - 10 µg of target antibodies and 2μg of spike-in antibody (anti-H2Av, Active Motif, #61686) were added and incubated on a rotator at 4°C overnight. ChIP samples were added with 50μL prewashed protein A Dynabeads (ThermoFisher) and incubated for 3h at 4°C. Beads were then collected on a magnetic rack and washed twice with 1mL cold RIPA buffer, twice with 1mL cold RIPA buffer containing 300mM NaCl, twice with 1mL cold LiCl buffer and once with PBS. Beads were then eluted with 100 μL buffer containing 0.1M NaHCO_3_, 1% SDS, and 20μg Proteinase K at 65°C overnight. Input sample volumes were adjusted to 100μL with the elution buffer. Samples were then purified using QIAquick PCR purification kit (Qiagen) and eluted in 30 μL 10mM Tris-HCl. For ChIP-Seq, entire ChIPed DNA or 300ng of the input DNA were used to construct libraries using NEBNext^®^ Ultra™ II DNA Library Prep kit with AMPure XP magnetic beads (Beckman Coulter). Library quality and quantity were estimated with Bioanalyzer and Qubit assays. The final libraries were sequenced on Illumina HiSeq 3000.

### ATAC-Seq library preparation

The Assay for Transposase-Accessible Chromatin with high-throughput sequencing (ATAC-Seq) was performed as described^48^. For each ATAC reaction, briefly, 5 × 10^4^ freshly collected cells were aliquoted into a new tube and spun down at 500 × *g* for 5 min at 4 °C. The cell pellet was incubated in 50 µL of ATAC-RSB buffer (10 mM Tris-HCl pH 7.4, 10 mM NaCl, 3 mM MgCl_2_) containing 0.1% NP-40, 0.1% Tween-20, and 0.01% digitonin (Promega) on ice for 3 min and was washed out with 1 mL of ATAC-RSB buffer containing 0.1% Tween-20. Nuclei were collected by centrifugation at 500 × *g* for 10 min at 4 °C, then resuspended in 50 µL of transposition reaction buffer containing 25 µL 2x Tagment DNA buffer, 2.5 µL transposase (100 nM final; Illumina), 16.5 µL PBS, 0.5 µL 1% digitonin, 0.5 µL 10% Tween-20, and 5 µL H_2_O. The reaction was incubated for 30 min at 37 °C with mixing (1000 r.p.m.) and directly subjected to DNA purification using the MinElute Reaction Cleanup Kit (Qiagen) according to the manufacturer’s instructions. Eluted DNA was amplified with PCR using Nextera i7- and i5-index primers (Illumina). Purification and size selection of the amplified DNA were carried out with AMPure XP magnetic beads (Beckman Coulter) to remove primer dimers and >1,000 bp fragments. For purification, the ratio of sample to beads was set to 1:1.8, whereas for size selection, the ratio was set to 1:0.55. Sequencing libraries were analyzed with Qubit and sequenced on HiSeq3000.

### Computational analysis

#### RNA-Seq data analysis

Raw sequencing data were aligned to the mouse genome mm9 using STAR software^49^. Reads on exons were collected to calculate reads per kilobase per million (RPKM) as a measure of gene expression level. Only genes with exonic reads of RPKM > 1 were considered expressed. For comparing gene expression levels before (D-3) and during (D2) adipogenesis, or in control and SMARCB1-deficient cells (Fig 2-S1), fold change cutoff of > 2.5 was used to identify differentially expressed genes. Gene ontology (GO) analysis was done using DAVID with the whole mouse genome as background (https://david.ncifcrf.gov).

#### ChIP-Seq peak calling

Raw sequencing data were aligned to the mouse genome mm9 and the drosophila genome dm6 using Bowtie2 (v2.3.2)^50^. To identify ChIP-enriched regions, SICER (v1.1) was used^51^. For ChIP-Seq of histone modifications (H3K4me1 and H3K27ac), the window size of 200 bp, the gap size of 200 bp, and the false discovery rate (FDR) threshold of 10^−3^ were used. For ChIP-Seq of non-histone factors, the window size of 50 bp, the gap size of 50 bp, and the FDR threshold of 10^−10^ were used.

#### Genomic distribution of ChIP-Seq peaks

To define regulatory regions (Fig 1C), combination of genomic coordinates and histone modification ChIP-Seq data were used. Promoter was defined as transcription start sites ± 1 kb. Promoter-distal regions were further separated into active enhancers (H3K4me1^+^ H3K27ac^+^), primed enhancers (H3K4me1^+^ H3K27ac^-^), and the other (H3K4me1^-^).

#### Motif analysis

To find enriched TF motifs in given genomic regions identified by ChIP-Seq, we utilized the SeqPos motif tool in the Cistrome toolbox (http://cistrome.org/ap/)^52^. For ChIP-Seq data with more than 5,000 regions (Fig 1, 3, 3-S1), top 3,000 significant regions were used. For ChIP-Seq data with small numbers (Fig 6 and 7), all regions were used. For differential motif analysis of given two regions (Fig 3-S2), HOMER software was used (http://homer.ucsd.edu/homer/)^53^.

#### Heat maps and box plots

The heat map matrices were generated using in-house scripts with 50 bp resolution and visualized in R using gplots package. Enhancers shown in the heat maps were ranked according to the intensity of SMARCA4 at the center of 400 bp window in control cells (Fig 4 and 5) or that of C/EBPβ in control cells expressing ectopic C/EBPβ (Fig 6 and 7). The ratio of normalized ChIP-Seq read counts in SMARCB1-deficient and control cells (Fig 4E and 5D), those in MLL4-deficient and control cells (Fig 5B), or those in SMARCA4-depleted and control cells (Fig 5F) in base 2 logarithm were plotted using box plot, with outliers not shown. Wilcoxon signed rank test (one-sided) was used to determine statistical differences in SMARCB1-deficient and control cells (Fig 4E and 5D), in MLL4-deficient and control cells (Fig 5B), or in SMARCA4-depleted and control cells (Fig 5F).

#### ATAC-Seq data processing

Raw ATAC-Seq reads were processed using Kudaje lab’s ataqc pipelines (https://github.com/kundajelab/atac_dnase_pipelines) which included adapter trimming, aligning to mouse mm9 genome by Bowtie2, and peak calling by MACS2. For downstream analysis, we used filtered reads that were retained after removing unmapped reads, duplicates and mitochondrial reads.

#### C/EBPβ-activated enhancers

To identify C/EBPβ-activated enhancers in preadipocytes (Fig 6 and 7), we first selected C/EBPβ^+^ AEs (C/EBPβ^+^ H3K27ac^+^ promoter-distal regions) in ectopic C/EBPβ-expressed cells. We further narrowed the focus down to SMARCA4^+^ ARID1A^+^ MLL4^+^ ones to better assess direct relationship between BAF and MLL4. We compared H3K27ac enrichment intensities in control and ectopic C/EBPβ-expressed cells. Enhancers with over 2-fold increased H3K27ac intensities in ectopic C/EBPβ-expressed cells were considered as C/EBPβ-activated enhancers.

## Accession numbers

All datasets described in this paper have been deposited in NCBI Gene Expression Omnibus under access #GSE151115.

## Acknowledgement

We thank Keji Zhao for *Smarca4*^AID/AID^ mice, Charles W. M. Roberts for *Smarcb1*^f/f^ mice, Ruth Kopyto for assistance in schematic drawing, Guojia Xie for suggestions, NIDDK Genomics Core and NHLBI DNA Sequencing and Genomics Core for next generation sequencing. This work was supported by the Intramural Research Program of NIDDK, NIH to K.G.

## Author Contributions

Conceptualization, Y.-K.P., J.-E.L., and K.G.; Methodology, Y.-K.P., J.-E.L., Z.Y., W.P., and W.W.; Investigation, Y.-K.P., T. O., and K.M.; Software, Formal Analysis, and Data Curation, J.-E.L. and W.P.; Writing – Original Draft, Y.-K.P., J.-E.L., and K.M.; Writing – Review & Editing, Y.-K.P., J.-E.L., K.M., and K.G.; Project Administration and Funding Acquisition, K.G.

## Declaration of Interests

The authors declare no competing interests.

## Figure legends

**Figure 1-Supple 1.**
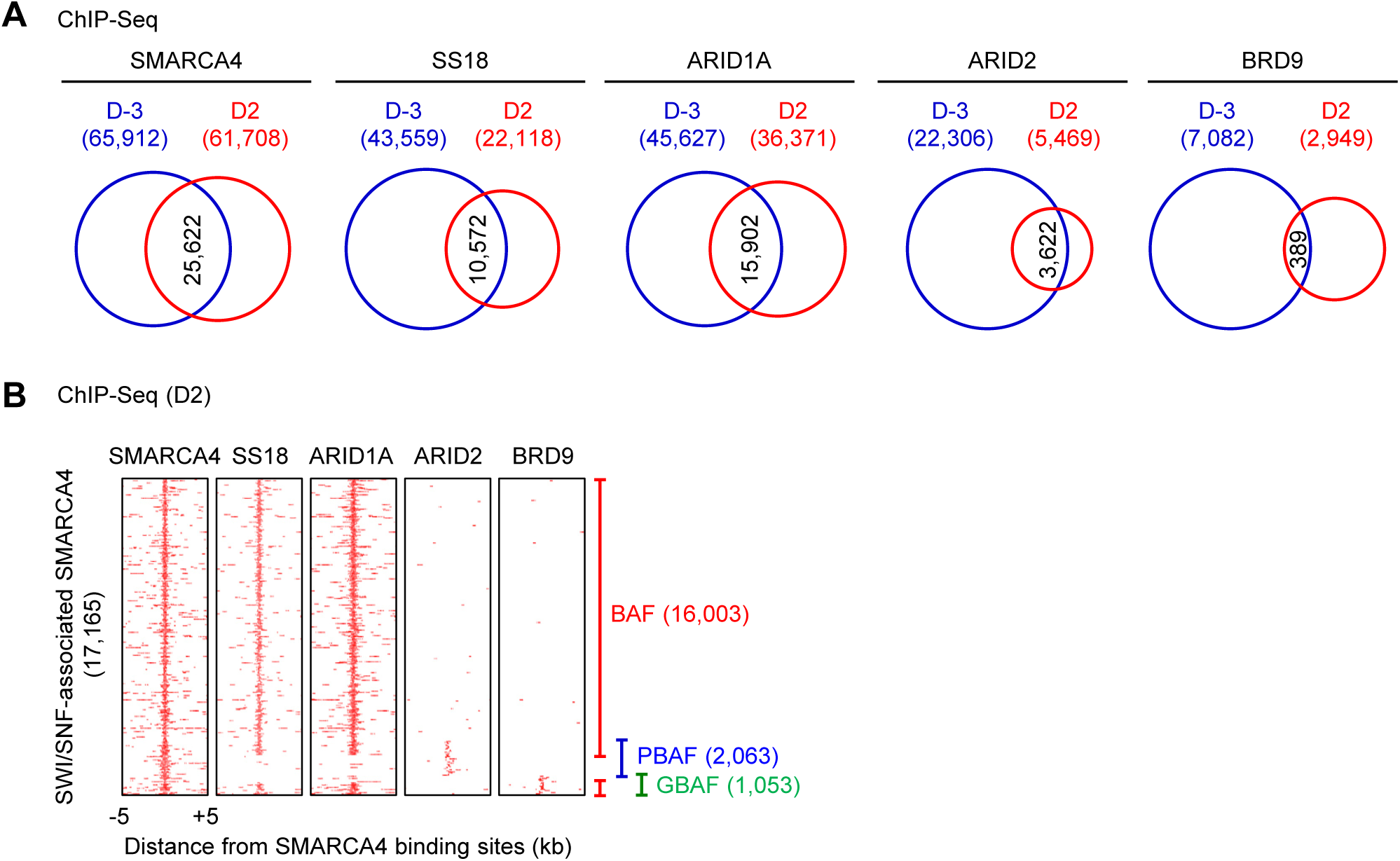
Genomic binding of SWI/SNF subunits in adipogenesis. **(A)** Venn diagrams depicting genomic binding of SMARCA4, SS18, BAF-specific ARID1A, PBAF-specific ARID2 and GBAF-specific BRD9 before (D-3) and during (D2) adipogenesis. Numbers of binding sites are indicated. **(B)** Heat maps for genomic binding of SMARCA4, SS18, ARID1A, ARID2 and BRD9 are shown around SMARCA4 binding sites at D2 of adipogenesis. Regions associated with either BAF, PBAF or GBAF are shown.

**Figure 2-Supple 1.**
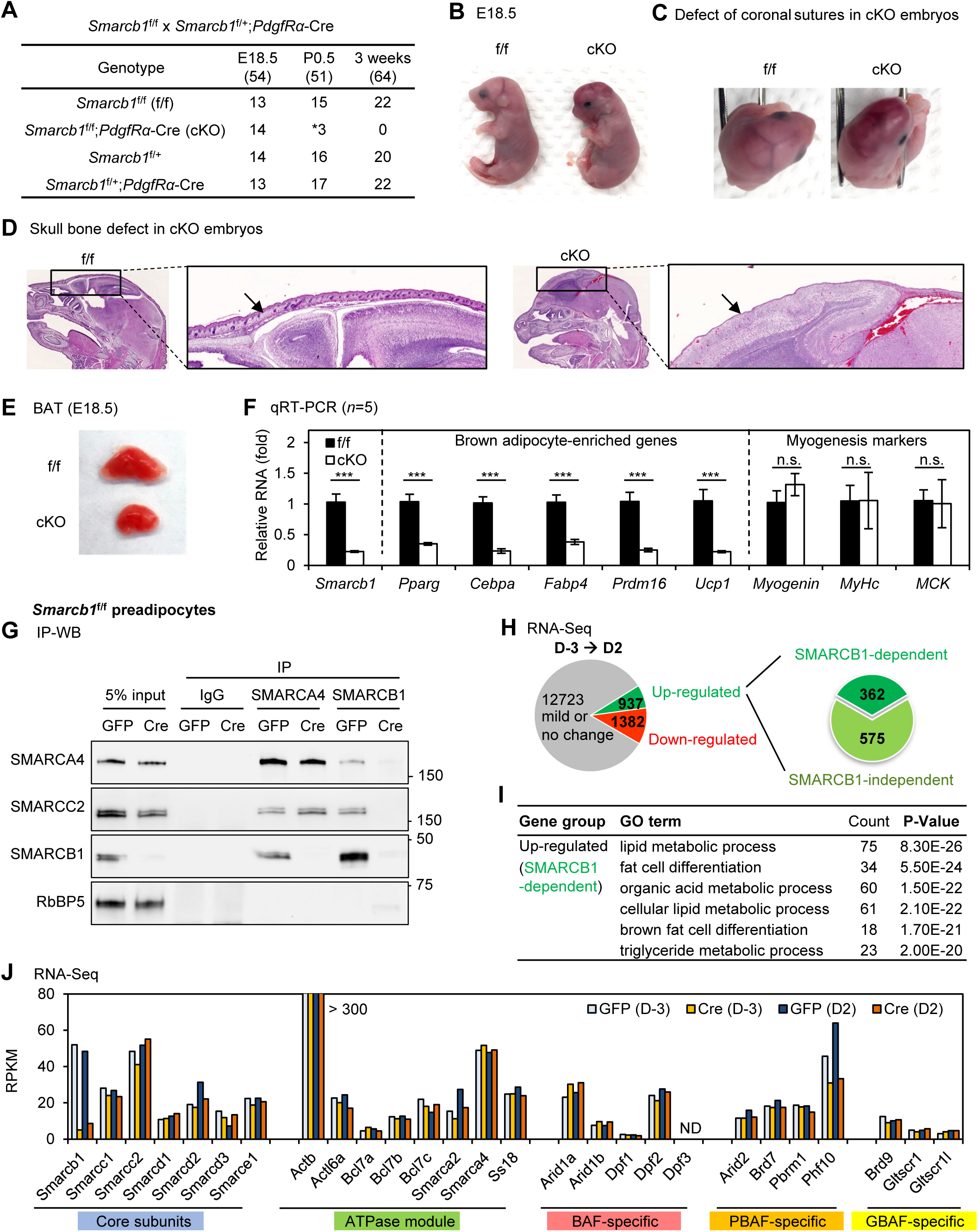
SMARCB1 is required for adipogenesis *in vivo* and in culture. (**A-D**) *Smarcb1*^f/f^;*PdgfRα-Cre* (cKO) mice showed defects in skull bone and died shortly after birth. **(A)** Genotype of progeny from the crossing between *Smarcb1*^f/f^ (f/f) and *Smarcb1*^f/+^;*PdgfRα-Cre* mice at embryonic day 18.5 (E18.5), post-natal day 0.5 (P0.5) and weaning (3 weeks of age). Dead pups are indicated by an asterisk. **(B)** Representative morphology of E18.5 embryos. **(C)** Defective coronal sutures in cKO mice. **(D)** H&E staining of the sagittal sections of head. Arrows indicate the skull bone. (**E-F**) Impaired BAT development in cKO mice. **(E)** Representative pictures of interscapular BAT. **(F)** qRT-PCR analysis of *Smarcb1*, adipocyte-enriched genes *Pparg, Cebpa, Fabp4, Prdm16* and *Ucp1* as well as myogenesis marker genes *Myogenin, MyHC* and *MCK* in BAT isolated from E18.5 f/f (*n* = 5) and cKO (*n* = 5) embryos. Quantitative PCR data is presented as means ± SEM. *** p<0.001. n.s., not significant. (**G-J**) SMARCB1 is required for adipogenesis in culture. Immortalized *Smarcb1*^f/f^ preadipocytes were infected with adenoviral GFP or Cre, followed by adipogenesis assays. Cells were collected at D-3 and D2 of adipogenesis for RNA-Seq. **(G)** Deletion of *Smarcb1* does not affect the interaction between SMARCA4 and SMARCC2. Nuclear extracts prepared at D2 of adipogenesis were immunoprecipitated (IP) with SMARCA4 or SMARCB1 antibody. Immunoprecipitates were analyzed by Western blot using antibodies indicated on the left. **(H)** RNA-Seq analysis at D2 of adipogenesis. Pie chart depicts SMARCB1-dependent and -independent up-regulated genes as well as down-regulated genes from D-3 to D2 of adipogenesis. The cut-off for differential expression is 2.5-fold. **(I)** Gene ontology (GO) analysis of 362 SMARCB1-dependent up-regulated genes defined in (**H**). **(J)** Deletion of *Smarcb1* does not affect the expression of other SWI/SNF subunits in adipogenesis.

**Figure 2-Supple 2.**
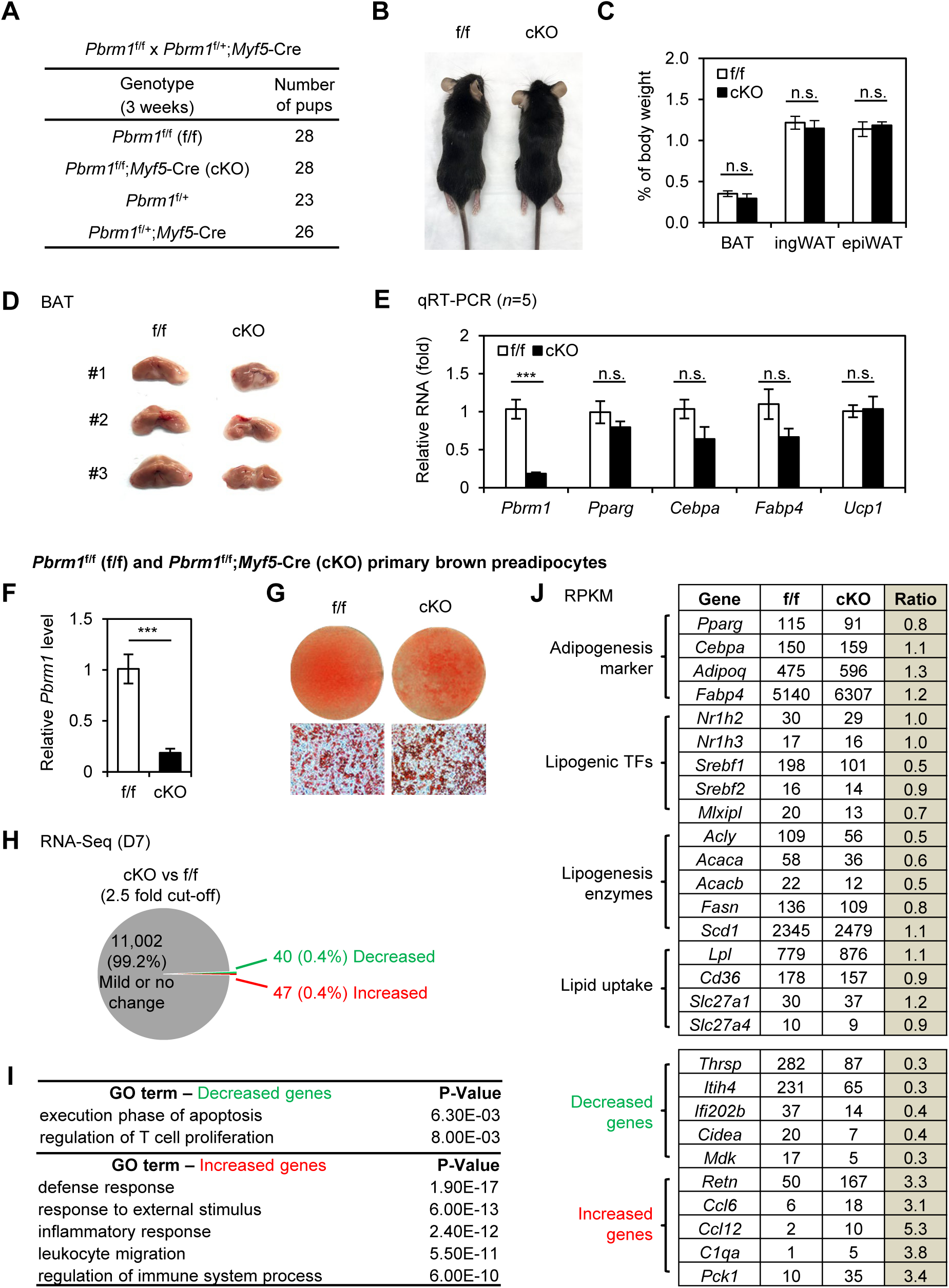
PBAF-specific subunit PBRM1 is dispensable for adipogenesis *in vivo* and in culture. (**A-E**) Characterization of adult (8 week-old) *Pbrm1*^f/f^;*Myf5-Cre* (cKO) mice. *Pbrm1*^f/f^ (f/f) were crossed with *Pbrm1*^f/+^;*Myf5-Cre* to obtain cKO mice. **(A)** Genotyping results. The expected ratios of the four genotypes are 1:1:1:1. **(B)** Representative pictures of 8-week-old f/f or cKO mice. **(C)** The average adipose tissue weights in f/f (*n* = 5) and cKO (*n* = 5) mice are shown as % of body weight. n.s., not significant. **(D)** Pictures of interscapular BAT isolated from three f/f or cKO mice. **(E)** Total RNA was extracted from BAT of f/f (*n* = 5) or cKO (*n* = 5) mice for qRT-PCR analysis of *Pbrm1*, adipogenesis markers *Pparg, Cebpα* and *Fabp4* and BAT marker *Ucp1*. Quantitative PCR data in all figures are presented as means ± SEM. *** p<0.001. n.s., not significant. (**F**-**J**) Pbrm1 is dispensable for adipogenesis in cell culture. Primary preadipocytes were isolated from f/f and cKO pups at P0.5, followed by adipogenesis until D7. **(F)** *Pbrm1* deletion in cKO preadipocytes was confirmed by qPCR of genomic DNA. *** p<0.001. **(G)** Oil Red O staining of differentiated cells. **(H)** RNA-Seq analysis of gene expression in differentiated f/f (*n* = 3) or cKO (*n* = 3) cells at D7. Pie chart depicts decreased or increased genes in cKO versus f/f cells. The cut-off is 2.5-fold. **(I)** Gene ontology (GO) analysis of decreased or increased genes defined in (**H**). **(J)** The list of representative adipocyte differentiation genes (upper panel) and significantly decreased or increased genes in *Pbrm1* KO cells (lower panel). RPKM values indicate gene expression levels.

**Figure 3-Supple 1.**
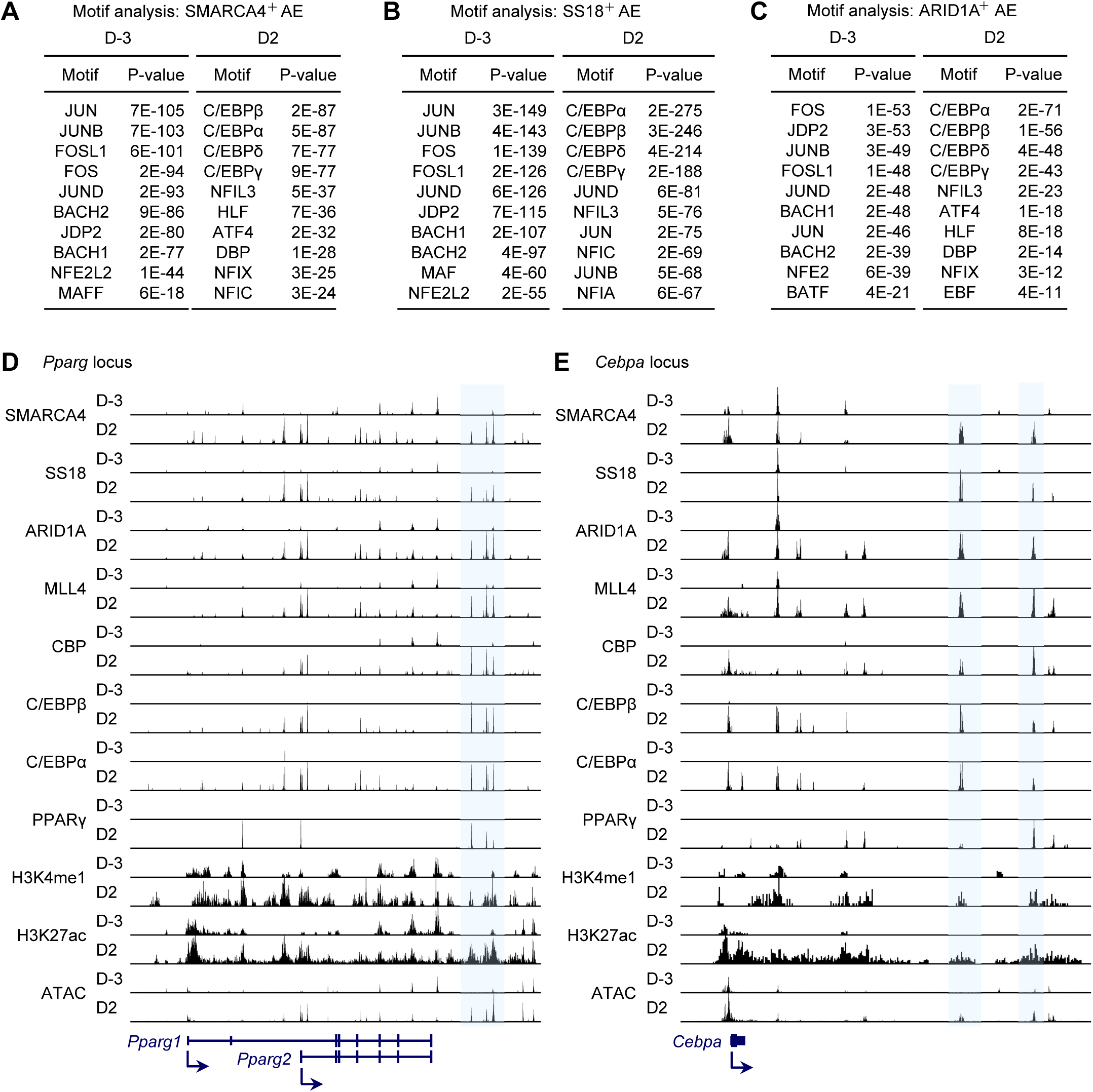
BAF co-localizes with LDTFs on active enhancers during adipogenesis. (**A-C**) Motif analysis of SMARCA4^+^ (**A**), SS18^+^ (**B**), or ARID1A^+^ (**C**) active enhancers (AEs) at D-3 and D2 of adipogenesis. Top 3,000 binding regions on AEs were used for motif analysis. (**D-E**) Genome browser shot of ChIP-Seq data of BAF subunits (SMARCA4, SS18 and ARID1A), MLL4, CBP, LDTFs (C/EBPβ, C/EBPα, and PPARγ), H3K4me1, and H3K27ac as well as ATAC-Seq data on AEs of *Pparg* (**D**) or *Cebpa* (**E**) gene loci.

**Figure 3-Supple 2.**
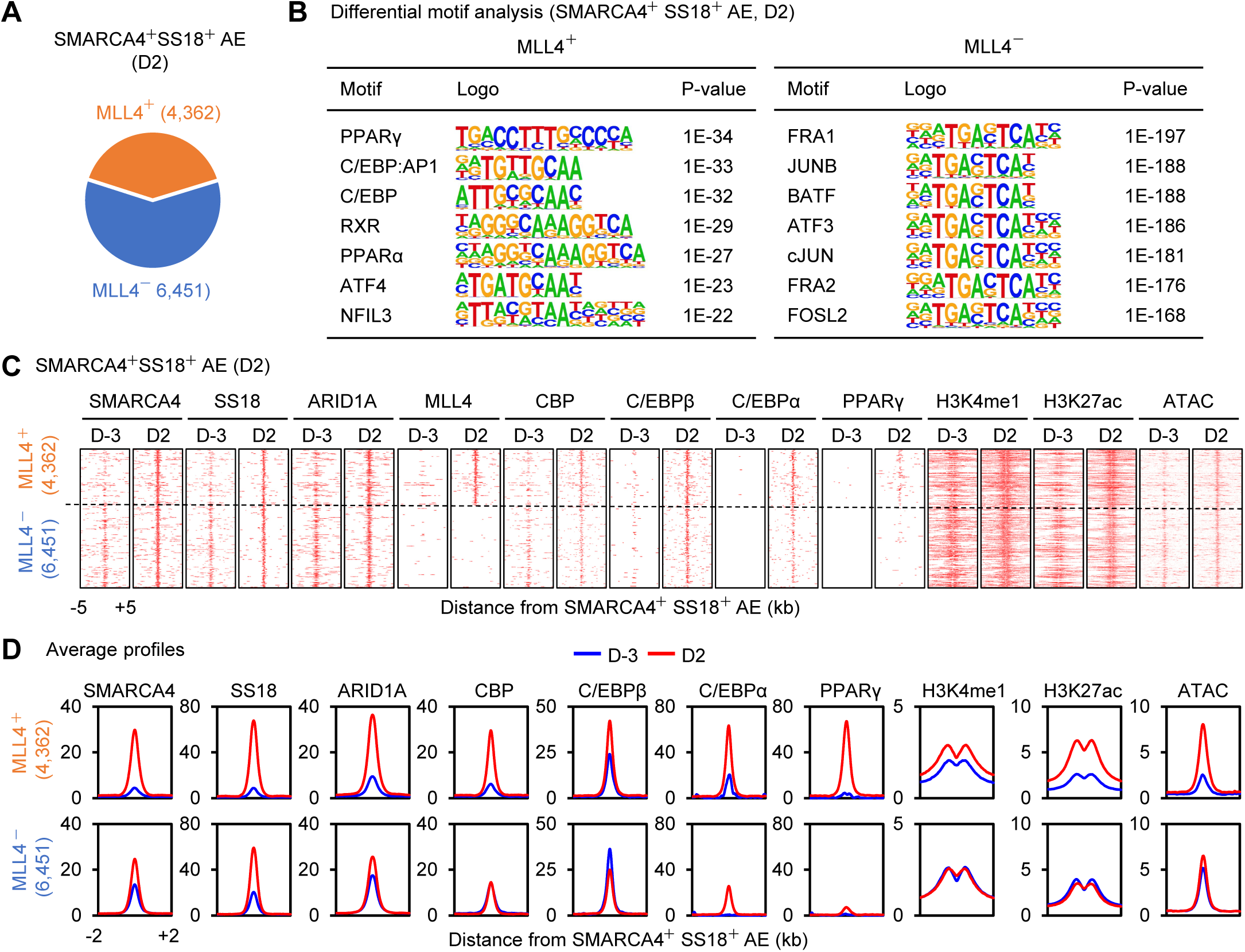
Markedly increased BAF binding and chromatin accessibility on MLL4^+^ active enhancers during adipogenesis. **(A)** Among the 10,813 BAF-binding (SMARCA4^+^ SS18^+^) active enhancers (AEs) at D2 of adipogenesis, 4,362 are bound by MLL4. **(B)** Differential motif analysis of MLL4^+^ or MLL4^-^ AEs at D2 of adipogenesis using HOMER. (**C-D**) Heat maps (**C**) and average profiles (**D**) around SMARCA4^+^ SS18^+^ AEs. Levels of CBP and H3K27ac were only induced on MLL4^+^ AEs. Normalized read counts are shown.

**Figure 3-Supple 3.**
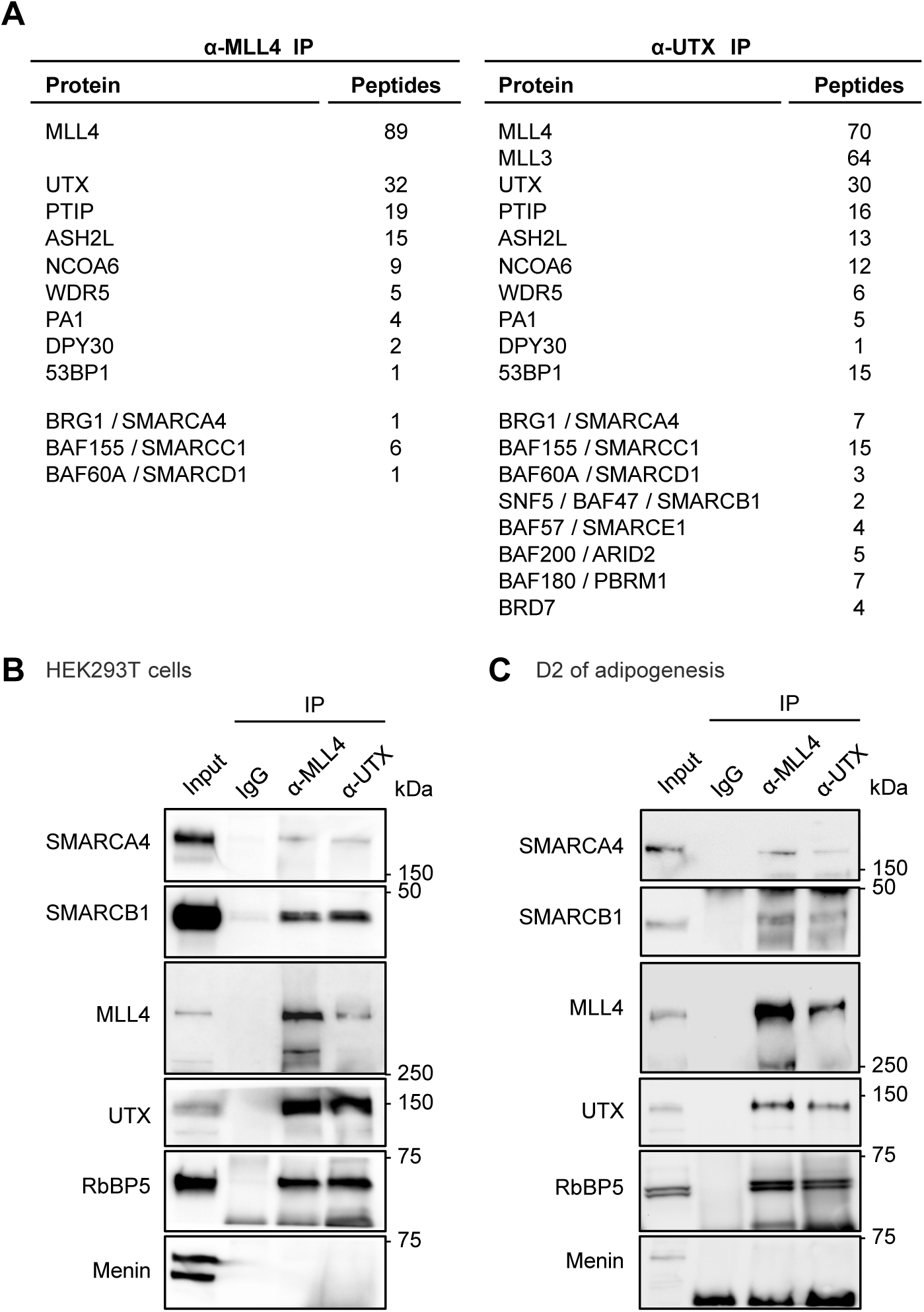
SWI/SNF subunits associate with MLL4 complex in cells. (**A**) MLL4- and UTX-associated proteins identified by IP-mass spectrometry analysis. Nuclear extracts prepared from mouse embryonic stem cells were subjected to immunoprecipitation with anti-MLL4 or anti-UTX antibody. Peptide numbers for IP-enriched BAF and PBAF complex subunits as well as MLL4 complex subunits are shown. (**B-C**) MLL4 complex associates with SMARCA4 and SMARCB1 in cells. Nuclear extracts from HEK293T cells (**B**) or preadipocytes during (D2) adipogenesis (**C**) were subjected to immunoprecipitation with anti-MLL4 or anti-UTX antibody. Immunoprecipitates were analyzed by Western blotting with antibodies indicated on the left.

**Figure 4-Supple 1.**
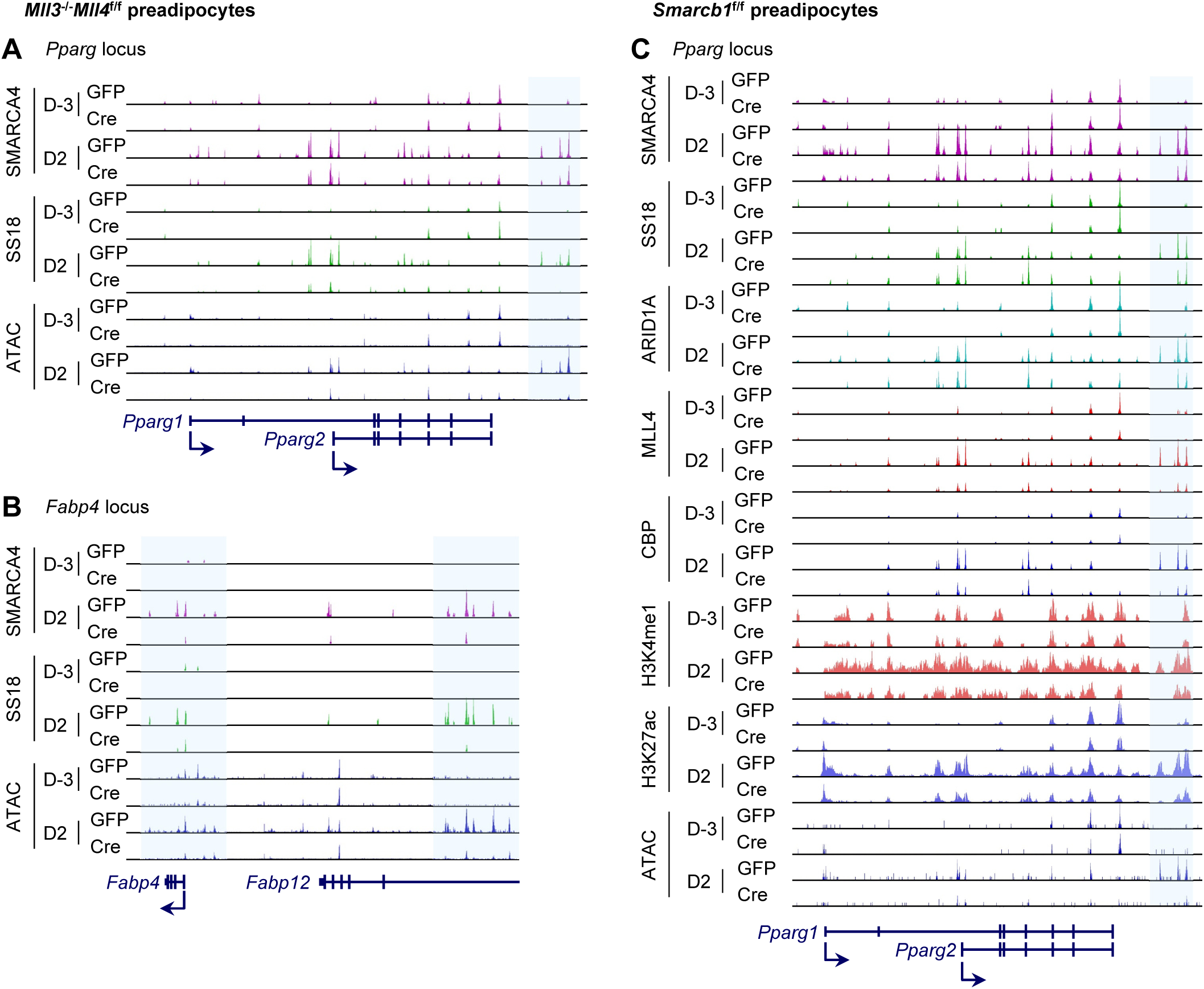
Reciprocal regulation between BAF and MLL4 on active enhancers during adipogenesis. (**A-B**) MLL4 is required for SMARCA4 and SS18 binding and chromatin opening on adipogenic enhancers around *Pparg* (**A**) and *Fabp4* (**B**) genes at D2 of adipogenesis. (**C**) SMARCB1 is required for SMARCA4, MLL4, and CBP binding and activation of enhancers around the *Pparg* gene at D2 of adipogenesis.

## Notes

### Competing Interest Statement

The authors have declared no competing interest.

